# SpY-C: Supervised Learning of Phosphopeptide Sequence Constraints Enables Global Prediction of SH2 Domain Binding

**DOI:** 10.64898/2026.08.18.744962

**Authors:** Alekhya Kandoor, Anny Carolline Silva Oliveira, Kazuya Machida, Blagoy Blagoev, Kristen M. Naegle

## Abstract

Tyrosine kinase signaling for cell development and homeostasis in multicelluar organisms and a major biochemical contribution is by driving interactions between phosphorylated tyrosines (pY) and SH2 domain containing proteins. This assembly is so important to driving cell outcomes that a wide variety of experimental and computational approaches have been used to understand which SH2-pY interactions occur, which still remains a challenge given the immensity (more than 45,000 pY and 120 SH2 domains in the human proteome). Based on biophysical constraints suggested by comprehensive contact mapping, here, we ask whether an approach might consider first asking if pY sequences conform to the shared rules of SH2 domain recognition by developing a classification approach that combines diverse training data. A wide range of validation suggests this approach, SpY-C, can classify pY sites as having the potential, or not, to be involved in SH2 domain interactions. We find that a relatively small set of representative SH2 binders, integrated from different experimental techniques, provides good classification. We use this classifier to annotate the human phosphoproteome and individual experiments, to explore the consequences of using super-SH2 domain reagents for pY enrichment, and to analyze the effects of mutations in altering pY site function. SpY-C provides a helpful step to more rapidly annotating pY function and for possibly improving machine learning approaches focused on specific SH2-pY interactions downstream of a first pass classification approach.

## Introduction

Cellular responses are coordinated through the dynamic assembly and disassembly of protein interaction networks in response to intra- and extracellular cues. Reversible tyrosine phosphorylation (pY) acts as one of the key regulatory switches underlying these processes by creating transient docking sites for signaling proteins [16]. A key component of this mechanism is the recognition of pY-containing motifs by Src Homology 2 (SH2) domains [40, 29], the primary phosphotyrosine-recognition modules that mediate phosphotyrosine-dependent protein–protein interactions, driving the assembly of signaling complexes and the propagation of signals regulating cell growth, differentiation, survival, and immune function [39, 34, 38, 41]. The human proteome encodes approximately 120 SH2 domains and contains more than 46,000 reported phosphotyrosine sites, representing a highly complex interaction landscape. However, systematically defining SH2–pY interactions remains experimentally challenging due to the enormous combinatorial search space and the dependence of binding specificity on both the SH2 domain and the surrounding peptide sequence. Several experimental approaches, including peptide arrays, affinity-based binding assays, structural studies, and phosphoproteomics, have substantially advanced our understanding of SH2-mediated recognition [43, 55, 9, 46, 18, 54, 50, 44]. Nevertheless, these approaches are limited in scale, often characterize only a subset of SH2 domains or pY peptides, and cannot comprehensively map the SH2 interaction landscape across the proteome. Consequently, although phosphoproteomic studies have identified tens of thousands of pY sites, only a small fraction have been linked to downstream SH2-mediated interactions, leaving much of the SH2 signaling network functionally uncharacterized. Computational methods have therefore emerged as an attractive strategy for predicting SH2–phosphotyrosine interactions. Existing approaches, including domain-specific predictors [22, 21], position-specific scoring matrix (PSSM)-based methods [36, 27, 54], energy-based models [25, 45], and structure-guided models [13, 47], have provided valuable insights into SH2 binding specificity. However, most are limited to individual SH2 domains or rely on position-wise scoring schemes that treat residues surrounding the phosphotyrosine independently, limiting their ability to capture context-dependent sequence determinants and reducing their applicability across the broader SH2 family. These experimental and computational challenges together underscore the need to develop methods that can identify SH2–phosphotyrosine interactions without requiring domain-specific models or exhaustive experimental characterization.

Our previous characterization of SH2–phosphotyrosine (pY) binding interfaces established a common structural framework for SH2 recognition. We found that peptide binding is primarily confined to residues spanning positions −2 to +4 relative to the phosphotyrosine, with residues outside this region making little direct contribution to the SH2 binding interface. Within this binding footprint, the Page 1 of 20 phosphotyrosine-binding pocket is highly conserved across the SH2 family and primarily accommodates the phosphorylated tyrosine together with immediate residues N-terminal and C-terminal to the pY, whereas the adjacent specificity pocket is considerably more variable and is largely responsible for peptide selectivity among SH2 subfamilies [19]. Consistent with these structural observations, extensive biochemical and structural studies have identified characteristic residue preferences within the C-terminal region flanking the phosphotyrosine for many SH2 subfamilies [52, 55, 3, 28]. These studies have defined canonical sequence motifs that capture the characteristic binding preferences of SH2 subfamilies. However, these models are largely based on independent positional preferences, assuming that individual residues contribute to binding in isolation. For example, an asparagine at the +2 position is a well-established determinant for GRB2 SH2 binding [42], yet experimentally validated GRB2-binding pY ligands lacking this canonical residue have been reported, indicating that deviations from preferred residues can reduce binding affinity without necessarily abolishing recognition [11]. Conversely, Pro or Leu at +3 position is a well-established determinant for Crk SH2 recognition, yet the presence of this canonical residue alone is insufficient for binding, as permissive residues at surrounding positions also contribute to recognition as shown in [30]. Together such observations suggest that SH2 recognition cannot be fully explained by single-position motifs alone, but instead depends on the broader sequence context and combinatorial relationships among residues within the binding interface. Additionally, the need to resolve coordinated binding of pY and flanking residues to the same SH2 residues suggest constraints that must be met for SH2 recognition. These findings led us to hypothesize that, despite the diversity of sequence preferences observed among SH2 subfamilies, SH2-binding pY peptides share common sequence features that reflect the conserved molecular mechanism of phosphotyrosine recognition. If such “rules” exist, it should be possible to learn these context-dependent features computationally and develop a global classifier capable of identifying SH2-binding phosphotyrosine sites independent of the specific SH2 domain.

Motivated by the conserved structural principles underlying SH2 recognition, we developed **Spy-C**, a global machine learning framework for predicting SH2-binding propensity from phosphotyrosine peptide sequence. We systematically evaluated multiple feature encoding strategies and training dataset compositions to identify a global representation that captures the shared determinants of SH2 recognition across this modular domain family. We validated the model using independent experimental datasets spanning multiple SH2 domains, diverse experimental platforms, and biological contexts to assess its robustness and generalizability. We then used this approach to annotate the human phosphoproteome, evaluate the effects of pY-enrichment strategies (comparing the engineered high affinity superbinder SH2 domain versus pY-antibody-based approaches), evaluated RTK-driven pY changes, and evaluated the effect of human mutations on altering the possible function of pY sites. The general learning framework is provided on GitHub, which has applications beyond SH2 domain recognition, the trained Python-based SpY-C model can be used to predict SH2 recognition for any sequence of interest, and the human phosphoproteome predictions are available for use in dataset annotation and protein visualization on ProteomeScout.

## Results

### Development of a global classifier to predict SH2–pY interactions

To test the central premise that binding rules exist, governed by the shared biophysical constraints of interactions, and they can be used to predict binding propensity, we first prototyped SVM modeling approaches on a small initial set of data. We used this preliminary dataset to determine model structure, including preferred feature encoding of the peptide sequences. Finding promising results, we then gathered diverse data across experimental approaches and with varying degree of SH2 domain and peptide coverage. We set aside specific validation sets and then sought to produce a final model that was trained and tuned well for generalizability across all of SH2 domain space based on cross-validation and validation. This general exploration identified interesting results regarding the depth and breadth of training data needed and we refer to the final trained model as SpY-C.

#### Feature encoding

Using a preliminary dataset, we evaluated multiple sequence and physicochemical feature encodings to determine which features best captured the characteristics distinguishing the positive and negative peptide sets. The preliminary dataset was built from Martyn et al. [32] affinity purification mass spectrometry (AP-MS), using pY peptides from K562 cell lysates from wild type SH2 interactions as positives and pY peptides from the same experiment found only by IMAC as negatives. Through this iterative feature-selection process, we found that a complementary combination of amino acid physicochemical properties (DPPS), hydrophobicity, and positional sequence preferences represented by log-odds scores provided the strongest separation between the two classes (Supplementary Figure c). Rather than retaining the full position-specific feature vectors, we found that collapsing each feature representation into an aggregated score substantially improved class separability while maintaining a direct connection to established biochemical determinants of SH2–pTyr recognition. This compact representation has also been used previously in the context of MHC–peptide binding prediction [10]. We subsequently used these features to train an initial support vector machine (SVM) model and evaluated its ability to distinguish the experimentally defined positive and negative peptide sets. The promising performance of this preliminary model demonstrated that a global predictive framework could be developed from experimentally derived SH2-binding data and, importantly, allowed us to identify and establish a compact set of sequence and physicochemical feature representations for subsequent model development.

#### Developing a diverse training dataset

We next focused on defining a comprehensive and representative set of peptides for model training. Although the K562 AP-MS dataset provided experimentally derived pTyr peptides enriched by wild-type SH2 domains, we recognized that the peptides recovered in individual experiments could be biased toward sequences that are more readily enriched or detected under the experiment conditions, rather than fully representing the broader diversity of sequence motifs capable of supporting SH2 domain binding. To address this limitation, we explored the PepspotDB dataset generated by Tinti et al. as a complementary source of positive training examples [50]. PepspotDB systematically characterized SH2 domain–pTyr peptide binding using a high-density peptide array platform comprising ~6,000 phosphotyrosine-containing peptides and profiling the binding specificity of ~65 human SH2 domains. Unfortunately, the database that housed PepspotDB is no longer available, and so we worked with the study authors to rebuild the resource from the original files, which we then translated to an updated phosphoproteome and used in this study for positive training data and for validation and testing.

To complement the expanded positive training set, we explored additional datasets to increase the number of negative examples. We initially incorporated the IMAC-only peptides from the K562 dataset as the negative training set. Because IMAC enrichment is driven primarily by the chemical affinity of phosphorylated peptides for the metal ions used in the enrichment and not sequence-based specificity, these peptides provide experimentally observed pY sequences that lack strong SH2-binding features. We further expanded the negative training set using another dataset from Chang et al., in which phosphotyrosine-containing peptides from pervanadate-treated HeLa cells were enriched in parallel using a SRC superbdinder (sSH2) domain or a cocktail of anti-pY antibodies, followed by IMAC purification [6]. From this dataset, we similarly selected peptides recovered only by IMAC enrichment and not by SH2-mediated enrichment. Together, this provided a total set of 205 negatives covering different pY sequence contexts from two cell lines.

#### Independent model validation data

To evaluate model performance independently of training, we assembled separate sets of experimentally characterized positive and negative peptides that were held out from model training. The positive evaluation set comprised several sources of known SH2-binding peptides, including established control peptides used in their SPOT array experiments [30], peptide sequences from PDB structures resolved in complex with SH2 domains, peptides found to bind wild-type SH2 domains in the K562 dataset, PepspotDB peptides detected across at least 4% of the SH2 domains tested and among the top 5% of signal intensities [50], and peptides experimentally confirmed to bind SH2 domains using Fluorescence Polarization assays [44]. Together, these datasets provided a diverse set of experimentally validated SH2-binding peptides for evaluating model sensitivity. For the negative evaluation set, we selected peptides from the HeLa dataset [6] that were recovered by either pY-antibody or IMAC enrichment but were not detected as binders by any type of the sSH2 domains tested. In total, the independent evaluation set contained 1,612 positive and 1,510 negative peptides. These held-out datasets were used to assess model sensitivity toward experimentally characterized known binders and specificity toward experimentally characterized high-confidence nonbinders, without overlap with the training data.

#### Exploring the best composition of positive training data across domains and measurement modalities

Having established the final feature encodings and a fixed set of experimentally derived negative set, we next focused on optimizing the composition of the positive training set. Unlike AP-MS enrichment, which identifies peptides that are recovered from a cellular phosphoproteome under specific experimental conditions, peptide SPOT array experiments directly test SH2 domain binding against a predefined set of peptides. We therefore compared training sets derived from AP-MS data, SPOT array data, and combinations of the two to determine which dataset or combination provided the strongest basis for building a global SH2-binding classifier. For this we developed an iterative framework in which candidate positive training sets were systematically constructed to assess three key aspects of the positive data: (i) the number of SH2 domains represented, (ii) the diversity and breadth of SH2-binding motifs captured by the selected peptides, and (iii) the experimental sources from which positive binding measurements were obtained. For each candidate positive training configuration, we trained an SVM classifier and evaluated its performance on the independent validation data (Figure 1A). This systematic comparison allowed us to determine how the composition of the positive training set influenced model generalizability and to identify the training configuration that provided the most robust balance between sensitivity and specificity (Figure 1A).

**Figure 1:**
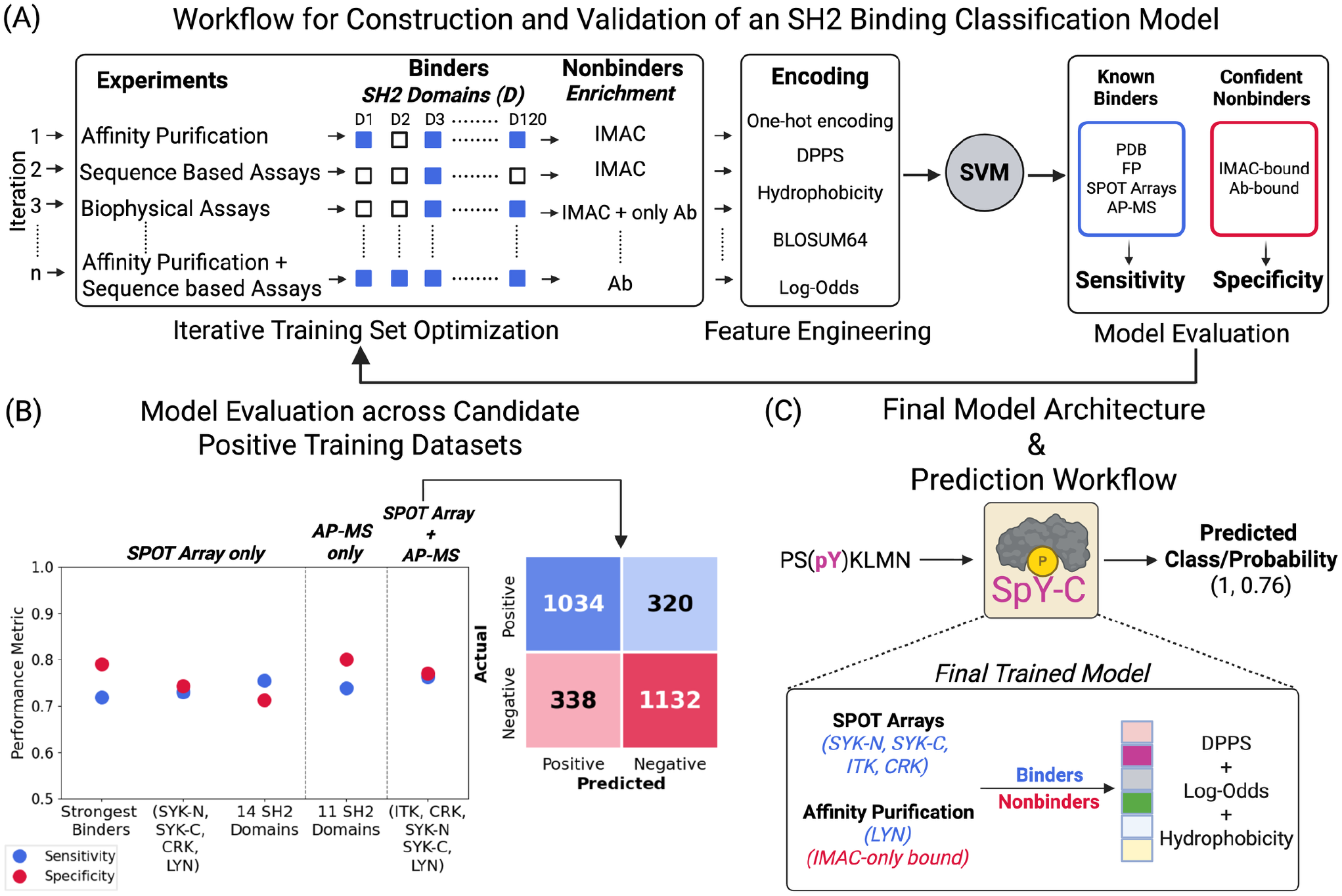
Development and implementation of a global SH2-binding classifier. **(A)** Overview of the iterative workflow used to develop a global classifier. We show some examples of how different combinations of experimentally derived datasets were evaluated to identify the optimal training strategy. Positive training sets were assembled from SH2 domain-bound peptides obtained from multiple experimental platforms, while negative training sets consisted of peptides with no evidence of SH2 binding. For each combination of training data, multiple sequence feature encodings were generated and evaluated within a support vector machine (SVM) framework. Model performance was assessed using independent evaluation datasets containing pY peptides that were never included during training. These evaluation datasets included experimentally validated SH2-binding peptides from diverse experimental methods, as well as confident nonbinders defined as those that failed to bind any wild-type or high-affinity SH2 domain variants in previous studies. Sensitivity and specificity on the independent datasets were used to compare model performance across iterations, allowing the training datasets and feature representations to be refined until the best-performing model was identified. **(B)** Candidate positive training datasets were evaluated while keeping the negative training set and sequence feature encoding fixed across all models. Model performance was assessed using a pooled independent evaluation set assembled from multiple experimental platforms. Sensitivity and specificity for each candidate training dataset are shown as scatter plots, including datasets generated from the strongest SH2 binders (top 5% of peptides bound by at least four SH2 domains), subsets of SH2 domains with limited or diverse binding specificities derived from the PepspotDB dataset (SPOT array-only), peptides bound by wild-type SH2 domains from the K562 dataset (AP-MS only), and the final training set comprising peptides integrated from multiple experimental sources (SPOT array + AP-MS). The confusion matrix for the final training dataset is shown on the right, summarizing the sensitivity and specificity achieved on the independent evaluation dataset. **(C)** SpY-C model used throughout this study. This final trained classifier was trained using experimentally validated binders from five SH2 domains compiled from SPOT peptide arrays and Affinity Purification (AP-MS), together with IMAC-derived peptides identified by AP-MS experiments as the negative training set. The final model combines log-odds scores, hydrophobicity, and DPPS sequence encodings within the SVM framework to generate a global predictor of SH2-binding propensity. Given a pY peptide sequence, SpY-C returns both a binding probability and a binary classification indicating whether the peptide is predicted to bind an SH2 domain, independent of the specific SH2 domain identity.

We first evaluated how the number of SH2 domains represented and the diversity of binding motifs captured influenced model performance. Using the PepspotDB dataset, we constructed candidate training sets by varying the stringency of peptide selection and the number of SH2 domains included. We first evaluated a set of the strongest-binding peptides and subsequently generated a more stringent subset comprising peptides within the top 5% of signal intensities and peptides that bound at least 4% of the SH2 domains tested. Although this stringent selection produced high specificity, it resulted in reduced sensitivity, indicating that restricting the positive set to the strongest or most broadly recognized binders limited the model’s ability to identify the full diversity of SH2-binding peptides. We next examined whether increasing the number of SH2 domains represented in the positive training set improved model performance. Specifically, we compared a smaller four-domain set comprising CRK, LYN, SYK-C, and SYK-N with a larger set representing 14 SH2 domains that included diverse binding specificities as laid out by Tinti et al. [50]. The model trained on the smaller four-domain set achieved a more balanced sensitivity and specificity than the model trained using the larger 14-domain set (Figure 1B). These results support the use of a smaller, carefully selected SH2-domain subset that captures a broader range of SH2-binding sequence preferences maintains a balanced sensitivity and specificity, compared to a larger set spanning more SH2 domains.

We next asked whether incorporating peptide-binding measurements from complementary experimental sources could improve model performance. We first evaluated the K562 dataset, which contains measurements for 11 SH2 domains from a single experimental platform. Despite the diverse number of SH2 domains represented, the model trained on this dataset showed reduced sensitivity. We therefore constructed a positive training set that combined measurements from multiple experimental platforms while retaining a smaller number of SH2 domains. Specifically, we included AP-MS-derived peptides for LYN [17] and SPOT-array measurements for four SH2 domains from two independent sources i.e. SYK-N, SYK-C, ITK from PepspotDB, and CRK from [30]. This multi-platform training set achieved the best overall balance between sensitivity (0.76) and specificity (0.77), outperforming models trained using data from individual experimental sources (Figure 1B). Together, these results suggest that combining complementary experimental sources, rather than simply increasing the number of SH2 domains represented, provides a more effective strategy for capturing diverse SH2-binding sequence preferences and improving the generalizability of the classifier.

Having determined a small, but diverse set of SH2 domain training data from complementary experiments performs the best, we next assessed whether the composition of a five-SH2-domain training set could be further optimized by expanding the number of domains or changing the SH2 domains represented improved the model. To determine whether the model had reached a performance plateau, we performed two complementary analyses. First, we cumulatively added additional SH2 domains to the training set and evaluated the resulting models at each addition (forward feature selection). The progressive inclusion of additional domains did not result in significant improvement in either sensitivity or specificity, indicating that expanding the number of SH2 domains beyond the initial set of five did not provide additional predictive benefit (Supplementary Figure c). Second, we systematically replaced individual SH2 domains within the training set with alternative domains, thereby modifying ~20% of the positive training examples at each iteration. These substitutions likewise did not consistently improve model performance, suggesting that the predictive information contributed by the original five domains was not substantially enhanced by exchanging them for other SH2 domains (Supplementary Figure c). Together, these analyses showed that the five-SH2-domain training set, which combined positive data from both AP-MS and peptide-array experiments, provided the best overall performance. This final training set is heterogeneous in both experimental source and biological context, i.e. combining data from multiple experimental sources and including peptides without restricting to their physiological relevance. This broader diversity likely allows the model to capture a wider range of sequence features that support SH2 recognition, improving its ability to generalize across independently derived datasets.

#### Final SpY-C model

Together, the final training dataset consisted of 309 experimentally validated binders and 205 high confidence non-binders. Using this optimized training dataset and feature encoding approach, we developed SpY-C (SH2-pY Classifier), a support vector machine classifier with a radial basis function (RBF) kernel that predicts the SH2-binding propensity of phosphotyrosine-containing peptides (Figure 1C). Previous studies have shown that the primary determinants of SH2 recognition are encoded within residues spanning positions −2 to +4 relative to the phosphotyrosine site. Accordingly, SpY-C uses this six-residue, pY-centered sequence window as its predictive input. For each peptide, the model returns both a binding probability score and a binary classification based on the optimized decision threshold (Figure 1C). To assess model generalization during development, SpY-C was evaluated using a nested cross-validation framework, which estimates predictive performance on held-out data drawn from the same distribution as the training set. SpY-C achieved a mean Accuracy of 0.871 ± 0.027, an F1-score of 0.892 ± 0.022, and a ROC-AUC of 0.931 ± 0.020 across the 10 outer folds. Following model selection, the final SpY-C model was retrained on the complete training dataset using the selected hyperparameters (C = 0.5, γ = 1.0) and evaluated on the independent validation datasets comprising 2,824 peptides (1,354 positives and 1,470 negatives). On these independent datasets, SpY-C achieved a Sensitivity of 0.764, Specificity of 0.770, Accuracy of 0.767, an F1-score of 0.759, and a ROC-AUC of 0.849, demonstrating robust predictive performance on experimentally independent data and represents a rigorous measure of SpY-C’s generalization to previously unseen data. This final trained model is used for all subsequent downstream analyses in this work.

### Comprehensive validation of SpY-C predictions

SpY-C was developed as a global classifier of SH2-binding motifs, rather than a predictor of interactions for individual SH2 domains. We therefore evaluated its ability to generalize beyond the training data by validating the model at two complementary levels. First, we assessed its performance on SH2 domains that were not represented during training. Second, we evaluated model predictions using multiple independent experimental datasets generated using orthogonal techniques than those used in the training dataset.

#### Generalization across SH2 domains

To assess generalization across unseen SH2 domains, we analyzed the PepspotDB dataset, which contains binding measurements for 63 SH2 domains, including 58 domains that were excluded from model training. For each SH2 domain, we identified peptides with signal intensities in the top 5 percentile and determined the fraction predicted by SpY-C to be SH2 binders (Predicted Binder Proportion). Across all the 63 SH2 domains, the average predicted binder proportion was ~80% (Figure 2A). Thirty-five of the 63 domains (~56%) exhibited binder proportions above this average, while the remaining domains also maintained relatively high predicted binder proportions, generally around 70%. Despite having no prior exposure to the majority of these SH2 domains, SpY-C consistently identified a high proportion of experimentally enriched peptides as SH2 binders, demonstrating that the model captures sequence features that generalize across the broader SH2 domain family. Additionally, we saw no pattern, such as better accuracy, with respect to performance in the domains used in training versus those not in training, suggesting we avoided overtraining to the selection set. PepspotDB data contains peptide sequences that at the time were predicted to be phosphorylated, but are still not yet annotated as phosphorylated. We restricted analysis to currently annotated pY peptides in the set. Compared with the full dataset, the average predicted binder fraction decreased modestly from approximately 0.80 to 0.75 (~5%). Despite this reduction, most SH2 domains retained predicted binder fractions between 60-75% (Supplementary Figure c), indicating that a large proportion of physiological phosphotyrosine sites remain compatible with SH2 domain recognition.

**Figure 2:**
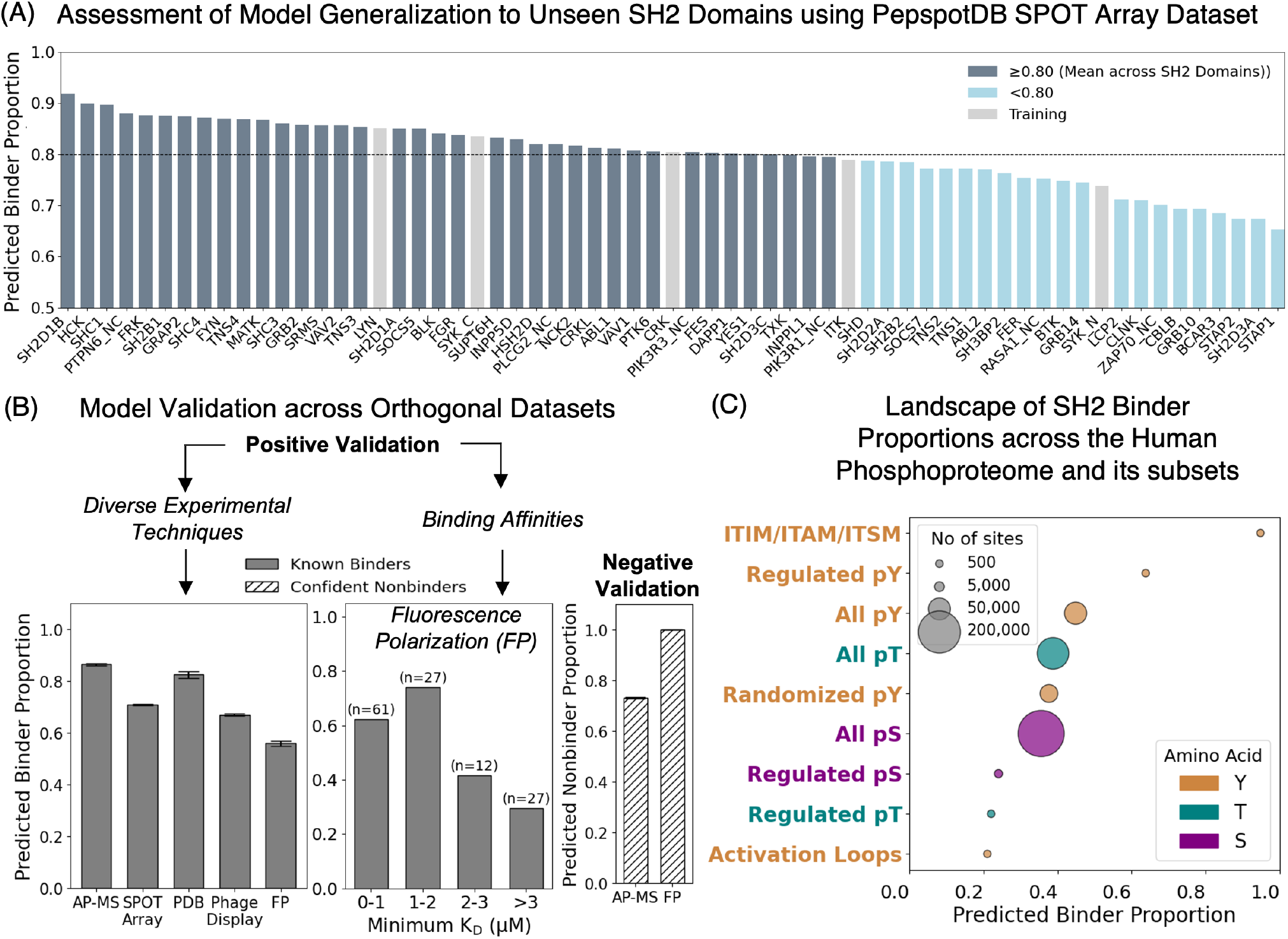
Validation of SpY-C predictions across SH2 domains, experimental platforms, and the human phosphoproteome. (A) Validation of SpY-C on the PepspotDB peptide array dataset [50], to assess SH2 domains not included during model training. For each domain, peptides within the top 5% of signal intensities were selected and the fraction predicted as SH2 binders by SpY-C was calculated (Predicted Binder Proportion). The mean binder proportion across domains was 0.80, with domains grouped above or below this value; domains included in model training are highlighted in grey. The consistent predictions across previously unseen SH2 domains support the generalizability of SpY-C across the SH2 family. **(B)** Positive and negative validation of SpY-C across multiple experimental techniques. Positive evaluation datasets included SH2-binding peptides identified from K562 dataset [32], all unique peptides within top 5% of each SH2 domain from PepspotDB dataset, SH2-bound peptides from PDB structures, pY peptides enriched by phage display [26], and fluorescence polarization (FP) binding assays [44]. FP peptides were further grouped by their strongest experimentally measured binding affinity (lowest K_D_), revealing a decrease in predicted binder proportion with increasing K_D_ (lowering affinity). Negative sets comprised IMAC- and pY-antibody-enriched peptides from [6] and FP peptides with no measurable binding to any tested SH2 domain [44]; the predicted nonbinder proportion is reported for each dataset. **(C)** Validation of SpY-C predictions across biologically relevant subsets of the human phosphoproteome. Predicted binder proportions were evaluated for established SH2-binding motifs (ITAM, ITIM, and ITSM) [31], activation loop pY sites from across 88 Human Tyrosine Kinases [33], and pS-, pT-, and pY-containing peptides from the human phosphoproteome. Additional sets included regulated phosphosites curated from PhosphoSitePlus and reported by Ochoa et al. [37], and a randomized pY control set generated by preserving pY position while randomizing the surrounding seven amino acids on N- and C-termini of pY sites located outside annotated protein domains.

#### Generalization across SH2 interactions found by different experimental techniques

We next evaluated SpY-C using independent datasets generated from diverse experimental approaches. To assess positive predictions, we compiled experimentally validated SH2-binding peptides from five orthogonal techniques and calculated the predicted binder proportion for each dataset. Peptides identified through K562 dataset and SH2-peptide complexes extracted from Protein Data Bank structures exhibited the highest predicted binder proportions (>80%), indicating excellent agreement between SpY-C predictions and experimentally observed SH2 interactions (Figure 2B). Intermediate binder proportions (~70%) were observed for peptides identified by PepspotDB and Phage Display experiments [26], both of which primarily characterize sequence-dependent SH2-binding preferences. The lowest overall binder proportion (~56%) was observed for peptides from biophysical assay such as Fluorescence Polarization (FP) [44]. (Figure 2B). To further investigate the FP dataset, we stratified peptides according to the lowest dissociation constant (K_D_) at which binding was experimentally detected, grouping peptides into increasing affinity ranges. Peptides exhibiting the strongest binding (lowest K_D_ values) showed the highest predicted binder proportions, with binder proportions progressively decreasing as K_D_ increased (Figure 2B). Although the overall predicted binder proportion across all unique FP binders was modest, this stratified analysis demonstrates that SpY-C preferentially identifies high-affinity SH2-binding peptides, suggesting that the high affinity interactions conform better to the rules of SH2 domain recognition.

We also assessed model performance on experimentally determined confident nonbinding peptides. Within the FP dataset, eight peptides showed no binding to any of the 80 SH2 domains tested experimentally, and SpY-C correctly classified all these as nonbinders (Figure 2B). As an additional negative validation, we analyzed phosphopeptides identified in the Chang et al. HeLa dataset [6] that were enriched by IMAC or antibodies, but were not recovered by any type of SH2 domain. Approximately 73% of these peptides were predicted as nonbinders by SpY-C, consistent with their lack of experimental SH2 binding. These complementary negative validation datasets demonstrates that SpY-C not only recognizes sequence features associated with SH2 binding but also accurately identifies phosphotyrosine peptides that are unlikely to function as SH2 ligands. This validation analysis shows that SpY-C generalizes effectively beyond the SH2 domains and datasets used for training. The model accurately identifies SH2-binding peptides across previously unseen SH2 domains and across multiple orthogonal experimental platforms while maintaining strong performance on experimentally validated nonbinding peptides. These findings support the robustness and broad applicability of SpY-C for predicting SH2-binding propensity across diverse biological and experimental contexts.

#### SpY-C recapitulates established SH2-binding biology and estimates SH2-binding potential across the human phosphoproteome

In addition to validating SpY-C using experimentally determined SH2-binding interactions, we sought to determine whether the model recapitulates established biological principles governing SH2-mediated signaling within the human phosphoproteome. Phosphotyrosine sites participate in diverse cellular functions, including enzymatic regulation, protein-protein interactions, and conformational control, but it is currently unknown what proportion of human pY sites serve as docking sites for SH2 domains. Hence, we deployed SpY-C to estimate SH2-binding propensity across broader phosphosite populations and use biologically well-characterized classes of phosphotyrosine sites with known SH2-binding behavior to provide an orthogonal validation of model predictions.

We first applied SpY-C to the curated human phosphotyrosine proteome and found that 45% of phosphotyrosine sites are predicted to function as SH2-binding motifs (Figure 2C). Because the complete set of SH2-binding phosphotyrosine sites in the human proteome is unknown, the accuracy of this global estimate for the pY population cannot be assessed directly. Instead, we evaluated whether the predicted binder proportion shifted in biologically expected directions across other control phosphosite subsets. First, to establish a sequence-based negative control, we generated a randomized phosphotyrosine dataset by shuffling residues flanking pY sites that are located outside structured protein domains, thereby disrupting the naturally occurring amino acid arrangement surrounding these pY sites. As expected, randomized phosphotyrosine sequences exhibited a lower predicted SH2-binding fraction than the native phosphotyrosine proteome (Figure 2C). The predicted binder proportion remained appreciable, however, likely because of sequence bias within the randomization window. Thus, while randomization does not eliminate all potential SH2-compatible motifs, the observed reduction indicates that SpY-C is sensitive to the disruption of evolutionarily selected sequence context. As a further control, we evaluated the sequences surrounding phosphoserine (pS) and phosphothreonine (pT) sites, treating them as if they central residue was a tyrosine phosphorylation and predicted using SpY-C. These phosphosite classes exhibited an even lower predicted SH2-binding fraction than the control pY population, indicating that the model identifies distinctly different sequences around pT and pS sites, consistent with the evolutionary background of SH2-pY binding (Figure 2C).

We next investigated whether phosphosites with established regulatory functions exhibit distinct SH2-binding propensities. To do this, we used the high-confidence regulatory phosphosite set curated from PhosphoSitePlus (PSP), which was used by Ochoa et al. as the training set for their machine learning framework to assign functional relevance scores across the human phosphoproteome and generate a functional landscape of phosphosite regulation [37]. Functionally regulated phosphotyrosine sites displayed a higher predicted SH2-binding fraction than the overall phosphotyrosine population, whereas functionally regulated phosphoserine and phosphothreonine sites exhibited lower predicted binder fractions than their respective background populations (Figure 2C). These results indicate that regulatory phosphotyrosine sites from this set are preferentially enriched for SH2-mediated signaling, while regulatory serine and threonine phosphorylation primarily participates in signaling pathways that are independent of SH2-domain recognition.

Together, these progressive shifts in predicted binder proportions demonstrate that SpY-C can distinguish biologically meaningful phosphotyrosine sequence contexts and provides confidence that the predictions across the human phosphotyrosine proteome reflects genuine signaling potential rather than classifier bias. Next, we sought to examine two classes of pY sites that might be expected to have differences in their ability to engage SH2 domains – immunoreceptor tyrosine-based inhibitory, switch, and activation motifs (ITIMs, ITSMs, and ITAMs), which are canonical SH2-binding motifs that mediate the recruitment of SH2-containing signaling proteins during immune receptor signaling and kinase activation loop pY sites, which we hypothesized would be less likely to bind SH2 domains as SH2 interaction with the activation loop might interfere with catalytic activity. Among 115 confidently annotated phosphorylated ITIM, ITSM, and ITAM sites from Liu et al. [31], SpY-C predicted ~95% as SH2 binders (Figure 2C), consistent with their established role as SH2-domain docking sites. In contrast, activation-loop phosphotyrosine sites showed the lowest predicted SH2-binding fraction among the evaluated phosphosite classes, with only ~21% classified as SH2 binders (Figure 2C). These results demonstrate that SpY-C captures functional differences among phosphotyrosine classes, identifying sites enriched for SH2-mediated interactions while distinguishing regulatory phosphotyrosines that operate through alternative mechanisms. Finally, from these analyses by extending predictions from experimentally selected peptides to endogenous phosphotyrosine sites, SpY-C is promising for capturing the sequence determinants underlying SH2-domain recognition and for estimating the fraction of pY sites across the human phosphoproteome that are likely to participate in SH2-dependent signaling.

#### SpY-C evaluation of phosphoproteomic experiments

SpY-C predictions across the entire phosphoproteome provides the first estimate of the proportion of pY sites that might function to recruit SH2 domains, at least based on sequence context. Next, we wished to use SpY-C to evaluate individual phosphoproteomic experiments, specifically to ask two key questions: 1) how might using SH2 superbinder (sSH2) reagents as the pY enrichment strategy reshape pY network coverage and 2) in receptor tyrosine kinase and kinase-linked receptor activation, what proportion of responsive pY sites function as SH2 recruitment sites. For this, we gathered a suite of experiments, focused on finding experiments where both traditional pY-antibody-based approaches and sSH2 enrichment were used and captured dynamic responses to receptor activation. As part of this, we complemented a prior pY-antibody-based enrichment of T Cell receptor (TCR) activation in Jurkat’s [8] with a new experiment, using super binder SRC sSH2 as an enrichment strategy.

#### SpY-C evaluation of pY-antibody-versus sSH2-enrichment

We examined whether the choice of affinity reagent used for phosphotyrosine peptide enrichment influences the sequence composition of the resulting phosphopeptide pool. To address this, we analyzed three independent studies representing distinct cellular contexts. The first dataset (HeLa) was generated by Chang et al. using the automated R2-pY workflow, in which phosphotyrosine peptides from pervanadate-treated HeLa cells were enriched in parallel using either the sSRC (SRC superbinder) or a cocktail of anti-pY antibodies, followed by IMAC purification [6]. The second dataset, from Bian et al., employed parallel sSRC superbinder and pY-antibody-based enrichment of phosphotyrosine peptides from Jurkat T cells [4] (referred to as Jurkat). Finally, our prior Chylek et al. study [8] performed pY-antibody-based enrichment in TCR-activated Jurkats, which we complemented with sSRC enrichment, allow us to evaluate specific signaling activation for comparison of enrichment strategy (referred to as Jurkat_TCR).

SH2 superbinders have previously been shown to provide greater depth of phosphotyrosine coverage than conventional pY-antibody-based enrichment. Consistent with these observations, analysis of the peptide pools used in this study showed that sSH2 enrichment recovered 1.7-fold and 2.1-fold more pY peptides than pY-antibody enrichment in the Jurkat and Jurkat_TCR datasets, respectively (Supplementary Figure c). We therefore asked whether this increased coverage simply expands the detectable pY landscape or preferentially enriches sites with intrinsic SH2-binding potential. Across all three datasets, SpY-C consistently predicted a greater proportion of SH2-binding phosphopeptides in superbinder-enriched samples than in pY-antibody-enriched samples (Figure 3A). Despite differences in cell type, sample preparation, and phosphoproteomic workflow, the direction and magnitude of the enrichment effect were remarkably consistent (7%), indicating that sSH2-based enrichment preferentially recovers phosphotyrosine peptides with sequence features favorable for SH2-domain recognition. We next asked whether this effect depended on the type of the engineered SH2 domain. Analysis of the Martyn et al. K562 dataset [32] showed similar predicted binder fractions across 11 independently engineered SH2 superbinders (Supplementary Figure c), suggesting that although individual superbinders differ in their sequence specificity and the breadth of peptides they capture, their recovered peptide pools are consistently enriched for general sequence features associated with SH2-domain recognition. To determine whether this selectivity limits recovery of particular classes of pY sites, we examined activation-loop phosphopeptides, which showed low overall SH2-binding potential in our analysis (Figure 2C). Although activation-loop peptides were occasionally recovered by sSH2 enrichment, all were predicted as nonbinders by SpY-C and were depleted relative to pY-antibody enrichment in both Jurkat datasets (Supplementary Figure c). These results demonstrate that, despite providing broader pY coverage, sSH2 enrichment remains selective for sequences compatible with SH2-domain recognition.

**Figure 3:**
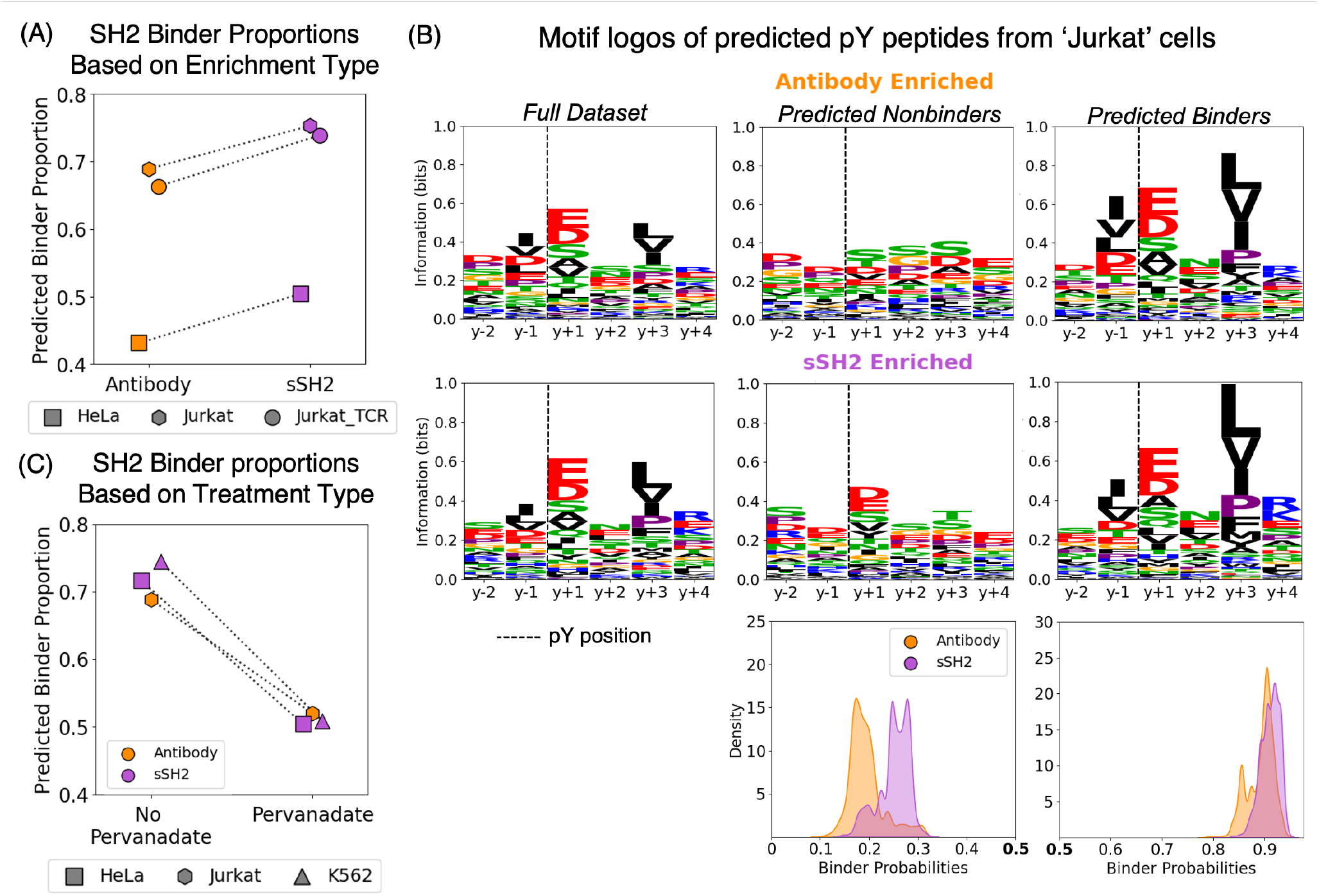
SpY-C reveals the impact of pY enrichment strategies on SH2-binding propensity. **(A)** We compared SpY-C-predicted binder proportions among peptides enriched by anti-pY antibodies and SH2 superbinders (sSH2) across phosphoproteomic datasets from HeLa [6], Jurkat [4], and Jurkat_TCR. For each dataset, we compared predicted binder proportions across matched enrichment strategies, experimental conditions, and cell lines. **(B)** Characterization of peptides predicted as nonbinders following pY-antibody- and sSH2-based enrichment using the Jurkat dataset [4]. Sequence logos were generated separately for peptides predicted as binders and nonbinders by SpY-C, with the phosphotyrosine (pY) position indicated by a dashed vertical line and excluded from motif generation. Predicted nonbinders recovered by sSH2 enrichment retained enrichment of acidic residues at the pY+1 position, a feature absent from pY-antibody-derived non-binders. To compare prediction probabilities between peptide populations, 50 unique peptides were repeatedly sampled without replacement, and the median SpY-C prediction probability was calculated for each sample over 5,000 iterations. Kernel density estimates (KDEs) of the resulting median prediction probability distributions are shown for the predicted binder and nonbinder populations from each enrichment strategy for the Jurkat dataset. The SpY-C decision boundary at 0.5 is indicated in bold.

Finally, we investigated the differences observed in the overall predicted SH2-binding distributions across datasets (Figure 3A), where the HeLa dataset exhibited a clear shift toward lower predicted SH2-binding propensity (Figure 3A). The HeLa dataset was unique, having used pervanadate treatment, prompting us to investigate whether phosphatase inhibition contributes to the observed shift in the predicted SH2-binding landscape. Hence, we evaluated three datasets that captured both pervanadate and non-pervanadate treated conditions (Chang et la. HeLa [6], Bian et al. Jurkat [4], and Martyn et al. K562 [32]), finding that, regardless of enrichment strategy, pervanadate drives an increasing proportion of phosphopeptides that do not match the rules of SH2 domain recognition (Figure 3C). Hence, this analysis highlights that SH2-binding potential of pY sites associated with expanded coverage introduced by phosphatase inhibition are less likely to function as SH2-domain docking motifs.

#### SpY-C reveals that sSH2 expand peptide recognition by relaxing canonical SH2 binding constraints

Superbinder SH2 domains were engineered by introducing mutations within the BC loop that enhance phosphotyrosine binding affinity while largely preserving the sequence specificity of their parental SH2 domains [20, 32]. Although these engineered domains retain the binding preferences the molecular basis underlying their expanded peptide repertoire remains poorly understood. Consistent with their intended function, sSH2-enriched datasets contained a substantially higher proportion of peptides predicted to bind native SH2 domains than pY-antibody-enriched datasets (Figure 3A). However, approximately 25% of peptides recovered by sSH2 enrichment were classified by SpY-C as non-binders. This observation raised an important question: do these peptides represent genuine non-binding sequences, or do they occupy a distinct region of sequence space that becomes accessible through the enhanced affinity of engineered sSH2 domains? We therefore used SpY-C to characterize the sequence features of sSH2-enriched peptides that are not predicted to bind native SH2 domains.

We first compared the sequence characteristics of predicted binders and predicted non-binders recovered from the Jurkat dataset (non-pervanadate conditions). Predicted binders from both enrichment strategies exhibited highly similar sequence profiles and closely recapitulate the characteristics of the training binder set (Supplementary Figure c). These sequence profiles highlight the expected enrichment of acidic residues at the +1 position and hydrophobic residues at the +3 position, relative to the pY site (Figure 3B) for SH2 binders. In contrast, predicted non-binders from both the groups were quite different. The pY-antibody-enriched dataset closely resembled the training non-binder set derived from IMAC-only phosphotyrosine peptide enrichment, suggesting that these peptides exhibit sequence characteristics similar to those observed in a phosphotyrosine-enriched background not selected through SH2-mediated enrichment. But a distinct pattern emerged among sSH2-enriched predicted non-binders, which retained acidic residue enrichment at +1 (Figure 3C). We assessed whether this pattern of +1 acidic enrichment was a general feature of sSH2 enrichment or a dataset-specific observation. Examination of additional sSH2-enriched datasets revealed a similar enrichment of acidic residues at the +1 position among predicted non-binders, with the strongest effect observed in the Jurkat_TCR dataset (Supplementary Figure c). Predicted non-binders consistently occupied a distinct region of sequence space characterized by retention of the +1 acidic feature. Importantly, these peptides did not resemble the training non-binder set (Supplementary Figure c), indicating that they were not simply representative of sequences lacking SH2-binding potential, but instead retained partial features associated with SH2 recognition.

To further resolve these peptide populations, we examined the continuous prediction probabilities generated by SpY-C. We used bootstrapping to build distributions of the median binding probabilities of predicted binders and non-binders from both enrichment types (Figure 3B). Predicted non-binders recovered by sSH2 enrichment exhibited consistently higher median prediction probabilities (~0.26) than pY-antibody-derived non-binders (~0.19), indicating that they were systematically more similar to the binder cutoff than pY-antibody-based non-binders. In contrast, predicted binders from both enrichment strategies exhibited similarly high prediction probabilities, with median values clustered near 0.9, although sSH2-derived binders showed a modest upward shift relative to pY-antibody-derived binders. The ability to explore sSH2 enrichment through SpY-C is a new look at exactly how the superbinders expand sequence space, which appears to extend breadth to sequences with a +1 acidic amino acid in particular.

#### Tyrosine phosphorylation-dependent signaling dynamically remodels the SH2-binding landscape in a receptor- and time-dependent manner

Having established that SpY-C accurately predicts SH2-binding phosphotyrosine ligands, we next sought to evaluate its utility in dissecting dynamic cell signaling. Phosphotyrosine-mediated signaling pathways are highly transient, with the temporal generation and removal of phosphotyrosine motifs regulating the recruitment of SH2-containing proteins to signaling complexes. Across diverse signaling systems, including receptor tyrosine kinase (RTK) and T-cell receptor (TCR) pathways, these dynamic phosphotyrosine-dependent interactions govern the assembly of signaling complexes and ultimately determine the specificity and duration of downstream signaling responses. We therefore analyzed time-resolved phosphoproteomic datasets to investigate how SH2-binding propensity evolves during signaling and to gain insight into the temporal organization of phosphotyrosine-dependent signaling networks.

#### SpY-C characterizes temporal binding shifts following receptor activation across species

To enable comparisons of SH2-binding dynamics across signaling systems, we assembled phosphotyrosine phospho-proteomic datasets (pY-antibody enriched) with comparable stimulation time points. We wished to gain a view across a range of RTK stimulation conditions and in so doing, aligned as best as possible the grouping of “early” and “late” dynamics and extended our predictions to the mouse proteome so we could evaluate a single dataset covering three RTKs with the same dynamics [2]. In all, we evaluated three human datasets: epidermal growth factor receptor (EGFR) at 1- and 8-minute post EGF [53], T-cell receptor (TCR) [8] at 1-minute post-stimulation, and vascular endothelial growth factor receptor (VEGFR) [56] at 10-minutes post VEGF-stimulation. The mouse dataset allowed us to evaluate PDGF, IGF, and FGF-stimulated responses at 3- and 15-minutes. Within each dataset, phosphotyrosine (pY) sites were classified as upregulated (log2 fold change ≥ 0.5) or downregulated (≥10% decrease relative to baseline) relative to unstimulated (0 min) controls at the selected early (1-3 minute) and late (8-15 minute) time points. The resulting upregulated and downregulated peptide groups were analyzed using SpY-C to predict SH2-binding potential, with predicted binder proportions quantified at each time point to assess temporal changes in SH2-binding potential across RTK stimulations and species (Figure 4A).

**Figure 4:**
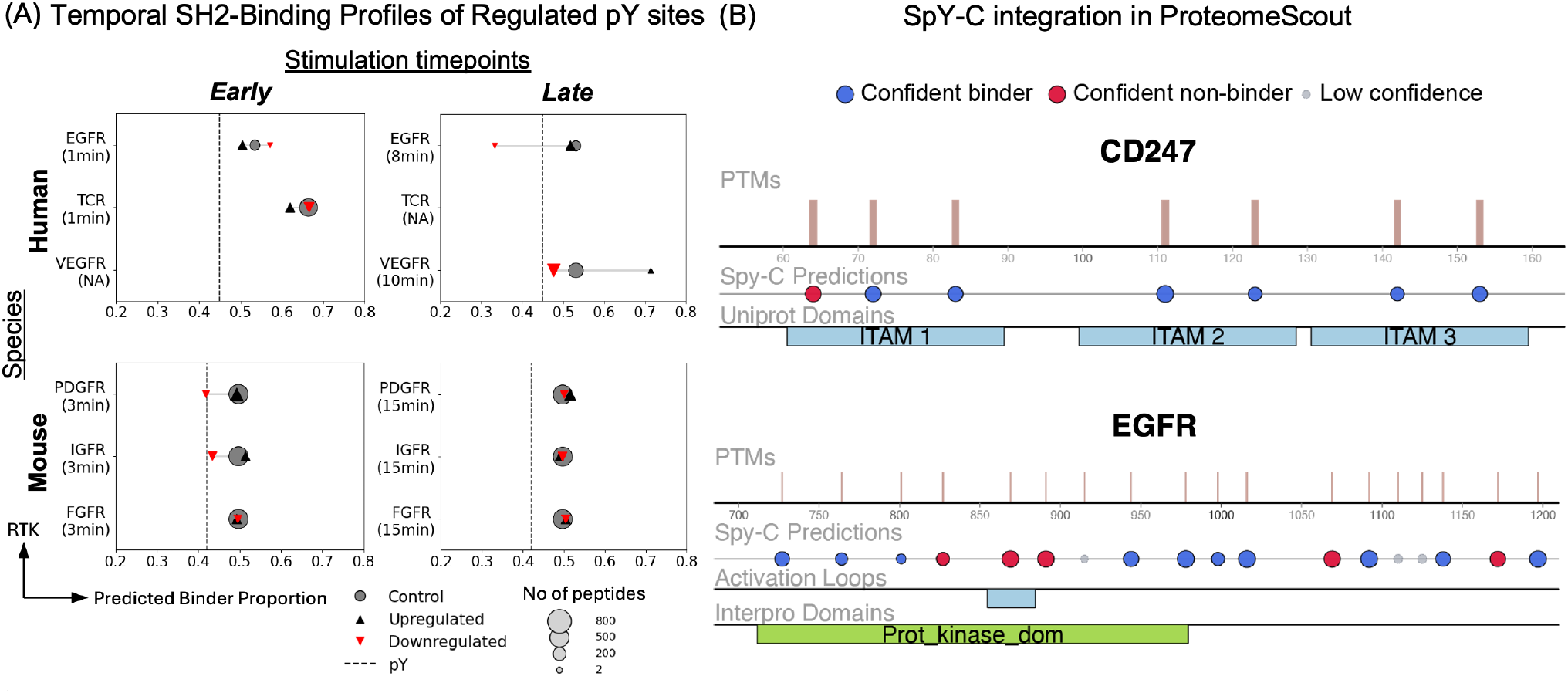
SH2-binding propensity of regulated pY sites during receptor tyrosine kinase signaling. **(A)** Predicted SH2-binding proportions of pY sites dynamically regulated following receptor tyrosine kinase (RTK) stimulation. For each phosphoproteomic time course in human and mouse cells, pY sites were classified as upregulated (log2 fold change ≥ 0.5) or downregulated (≥10% decrease in abundance relative to the corresponding control). SpY-C predicted binder proportions were calculated separately for the upregulated and downregulated pY sites at each time point. All pY sites identified across the complete time course were used as the corresponding experiment-specific control. Vertical reference lines indicate the predicted binder proportions for the pY sites from human and mouse reference phosphoproteomes. The size of each data point and the arrows markers are proportional to the number of pY sites within each regulated subset, providing context for the scale of phosphorylation changes observed during RTK signaling. **(B)** SpY-C predictions have been integrated into ProteomeScout for annotation and shown here, on the protein viewer. Example protein segments are shown for CD247 (CD28 epsilon) and EGFR. Circle sizes are proportional to the size of the probability of that category (large for high confidence in binder or non-binder) and predictions in the range near the SVM boundary are indicated as low-confidence. ProteomeScout protein viewer tracks were selected to indicate specific features, such as the location of pY in the kinase activation loop versus the C-terminal tail sites of EGFR.

Among the human datasets, receptor stimulation produced distinct temporal patterns of predicted SH2-binding potential. During very early signaling (1-minute), in both TCR and EGFR activation pY sites that immediately decreased in abundance showed higher predicted SH2-binding fractions than newly upregulated sites. At later time points (8-10 minutes), however, EGFR and VEGFR stimulation showed the opposite pattern, with upregulated pY sites exhibiting higher predicted SH2-binding proportions than downregulated sites (Figure 4A). In the mouse dataset, at 3-minutes following PDGFR and IGFR stimulation, upregulated pY sites exhibited higher predicted SH2-binding fractions than downregulated sites. By 15 min, these differences largely diminished, with upregulated and downregulated sites showing comparable SH2-binding proportions across all the receptors, suggesting convergence toward a more balanced phosphotyrosine signaling state (Figure 4A). FGFR was a notable exception, as upregulated and downregulated pY sites maintained similar SH2-binding propensity from 3 to 15 min, with both remaining close to the background control level. These results collectively suggest that phosphotyrosine sites generated during receptor stimulation are functionally heterogeneous, and their potential to engage SH2-containing proteins can change over time.

### Deployment of SpY-C predictions for general use

Given the relatively small degree of functional annotations on pY residues across the human proteome [35], the ability to classify pY sites as having the capacity or not to recruit SH2 domains, is useful. To this end, we integrated SpY-C predictions into an open source database of post-translational modifications, ProteomeScout [33]. In this integration, we indicate three total classes of SpY-C predictions: high confidence binders (prediction probabilities ≥ 0.6), high confidence non-binders (prediction probabilities ≤ 0.4), and low-confidence predictions that exist near the SVM boundary. These predictions are attached to human records in the ProteomeScout database file. Additionally, researchers can annotate a phosphoproteomic experiment with SpY-C predictions using the “Dataset Annotation” feature, such as was performed in the evaluation of RTK-specific binding events. Finally, the feature track in the protein viewer presents SpY-C predictions for pY sites. Figure 4B provides example protein sections covering SpY-C predictions on two receptors – CD247 and EGFR. Consistent with analysis in the ITAM/ITM/ITSM set, all known ITAM doublet sites show high probability of being recognized by SH2 domains. A third site in the first ITAM region is predicted to have low probability of interacting with an SH2 domain. In EGFR, consistent with the global activation loop analysis, the activation loop pY is predicted to have low probability of binding SH2 domains. As expected, the majority of C-terminal tail sites in EGFR are predicted to match SH2 domain binding rules, with Y1110 and Y1125 being near the classification boundary. However, two sites are confidently predicted to not recruit SH2 domains – Y1069 and Y1172. It is well established that both Y1069 and Y1172 drive interactions, with CBL and SHC, respectively. However, these protein interactions involve PTB (another phosphotyrosine binding domain), suggesting that, at least on EGFR, there is a strong separation in sequences that recruit SH2 domains, compared to those that recruit PTB domains, a potentially useful trait for increasing recruitment specificity and reducing competition.

### SpY-C predictions reveal key determinants of SH2–pY recognition and the consequences of their perturbation using known mutations

Beyond predicting SH2 binding interactions, we wished to evaluate the functional consequences of disease-associated sequence variation within known pY motifs and examined how these variants might alter SH2 domain recognition. This analysis provides an application for understanding how mutations may alter SH2 domain specificity and rewire signaling networks.

### Mapping disease associated mutations to known phosphotyrosine sites and predicting SH2 binding

We investigated the potential impact of disease-associated mutations on SH2-pY recognition by mapping the clinically annotated missense variants from ClinVar database [24] to known pY sites in the human phosphoproteome. This identified 6,387 missense mutations occurring within the canonical SH2 recognition window (pY-2 to pY+4), representing approximately 4% of all ClinVar missense variants (Figure 5A). These mutations were distributed relatively evenly across the six flanking positions, with each position accounting for ~16% of variants, although pY+1 showed a modest enrichment at ~19%. We also examined substitutions directly at the pY residue, where cysteine and histidine were the most frequent substitutions, both of which can arise from tyrosine through a single-nucleotide change. To assess the potential effects of mutations within the SH2-binding window on domain recognition, we used SpY-C to predict the binding potential of wild-type and mutant peptide sequences (excluding the central tyrosine mutations). The majority of variants (~85%) retained their predicted binding classification following mutation (Figure 5B), with predicted binders generally remaining binders and nonbinders remaining nonbinders (Figure 5B). The remaining 16% of variants altered their predicted classification, either resulting in a loss of binding (1→0: Binder to Nonbinder) or a gain of binding (0→1: Nonbinder to Binder).

**Figure 5:**
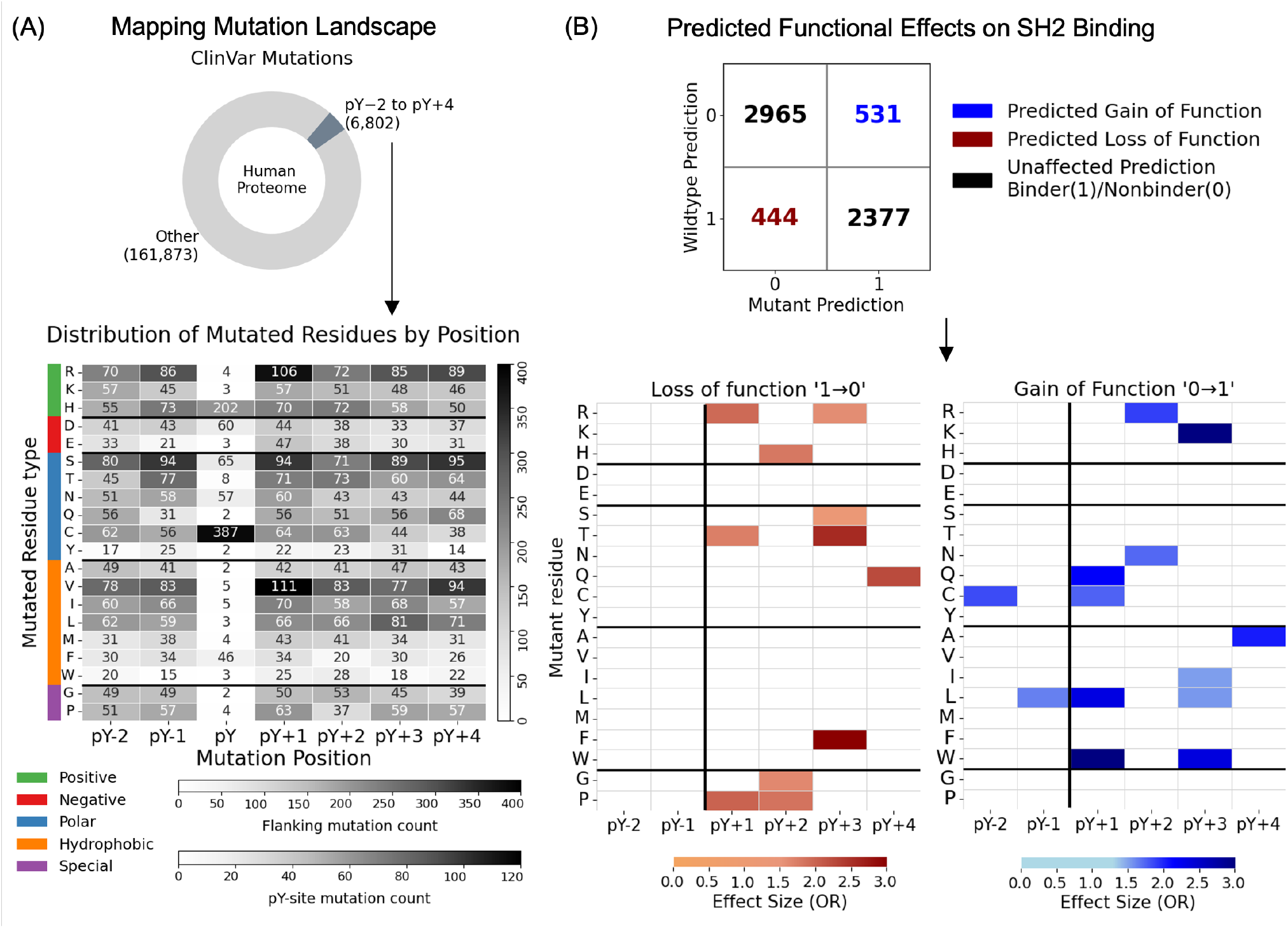
Predicting the impact of disease-associated mutations on SH2-binding using SpY-C. (A) Mapping of ClinVar disease-associated missense variants to pY peptides in the human phosphoproteome. Variants within the SH2 recognition window (pY-2 to pY+4) were retained, and the heatmap shows the distribution of mutated residues across positions. (B) Predicted effects of disease-associated mutations on SH2 binding. Wild-type–mutant pY pairs were classified based on changes in SpY-C predictions, with nonbinder-to-binder transitions defined as gain-of-function and binder-to-nonbinder transitions as loss-of-function. The confusion matrix summarizes preserved and altered binding predictions. Residue- and position-specific enrichment analysis identified substitutions associated with predicted gain- and loss-of-function events and only significant enrichments (FDR-adjusted P<0.05; odds ratio >1) are shown.

### Functional enrichment of mutations predicted to alter SH2 binding

We next examined whether SpY-C-classified wildtype-mutant pairs with altered binding predictions showed residue- and position-specific enrichment within the SH2 recognition window. For gain-of-function mutations (0→1), enrichment was assessed relative to peptides originating from the same initial prediction class (0→1 and 1→1), whereas loss-of-function mutations (1→0) were evaluated relative to peptides predicted to become non-binding (0→0 and 1→0).

Several consistent patterns emerged from this analysis. Mutations C-terminal to the phosphotyrosine, particularly at pY+1 to pY+3, accounted for most predicted gain- and loss-of-function events, consistent with the established role of these positions in SH2 specificity. No significant enrichment was observed at pY-2 or pY-1 among loss-of-function mutations, with only limited enrichment in the gain-of-function group (Figure 5B). At pY+3, loss-of-function pairs were enriched for substitutions to polar residues, particularly serine and threonine, whereas gain-of-function pairs showed strong enrichment for hydrophobic substitutions (Figure 5B). This is consistent with structural studies across protein-protein interaction interfaces showing that substitutions to smaller polar residues can disrupt favorable interactions within hydrophobic binding pockets, while even bulky residues such as phenylalanine may fail to preserve interactions when side-chain geometry differs [48]. Together, these patterns highlight pY+3 as a major determinant of mutation-induced changes in predicted SH2 binding.

We next identified enrichment patterns reflecting domain-specific SH2 preferences, despite SpY-C being designed as a global predictor of SH2-pY recognition. Predicted loss-of-function mutations recapitulated known non-permissive residues for the CRK SH2 domain, including histidine and glycine at pY+2, arginine at pY+1, and proline at pY+1 and pY+2 [30] (Figure 5B). Conversely, lysine substitutions at pY+3 were significantly enriched among predicted gain-of-function events. SUPT6H is the only SH2 domain in SMALI [15] reported to prefer lysine at this position, and a small number of binder peptides in our training set also contained pY+3 lysine. Thus, although uncommon, lysine at pY+3 can support SH2 binding in a favorable sequence context, illustrating how domain-specific preferences can emerge within the global SpY-C model. The recovery of these experimentally characterized sequence preferences provides additional context for interpreting mutation-induced changes predicted by the model. Finally, this demonstrates the applicability of SpY-C for systematically identifying and prioritizing disease-associated mutations that may alter SH2 binding for subsequent experimental investigation.

## Discussion

Our prior work, which involved the comprehensive extraction of SH2 domain interaction interfaces from structure, suggested, given the need to co-satisfy bonds between multiple ligand sites and the same SH2 residues, that there are shared rules peptides follow to bind to any SH2 domain. This led to a new approach to SH2 domain prediction – focused on classification of binding potential for pY-containing sequences to any SH2 domain versus first predicting whether a sequence will bind individual SH2 domains. Although there is no fundamental ground truth, we lack the experimental evidence at scale to truly evaluate all SpY-C predictions, the broad testing across withheld data, orthogonal data, and direct applications suggest that SpY-C predictions are biologically useful for stratifying pY peptide functions. This is broadly useful for multiple reasons: 1) it allows us to annotate possible functions to what has been a largely under annotated phosphoproteome and for contextualizing phosphoproteomic experiments and mutations and 2) a first pass classification might significantly improve the task for a secondary machine learning approach focused on predicting specific domain-pY interactions. However, a key caveat of these predictions is that they are entirely predicated on evaluating the match to necessary sequence requirements for SH2 domain recognition – it does not account for the linear accessibility in the broader protein context, which is also very likely important. Activation-loop pY sites illustrate this distinction: although these sites were generally less compatible with SH2 recognition, this does not preclude SH2 binding in specific contexts, as exemplified by SRC activation-loop pY recognition by the p85 N-SH2 domain when presented as an accessible linear phosphopeptide [51], or by recognition of the insulin receptor activation loop by the APS SH2 domain through a higher-order dimeric assembly [14]. Incorporating such structural accessibility and other cellular information into future iterations of the framework could help distinguish sequence-compatible interactions from those more likely to occur physiologically. An additional caveat for interpretation is that these predictions capture the function of pY in driving an interaction via an SH2 domain, but some pY sites might also drive interactions, just not by SH2 domains. In fact, in at least the few well annotated cases such as EGFR pY1173 a predicted nonbinder by SpY-C has been shown to support SHC recruitment through its PTB domain rather than its SH2 domain [12]. This suggests that the model may highlight largely orthogonal sequence requirements among phosphotyrosine reader domains such as SH2 and PTB, providing an interesting direction for further exploration.

The evaluation of sSH2 enrichment, compared to pY-antibody enrichment, was one orthogonal test of the fundamental approach – we expect that pY sites enriched by sSH2 domains should have a higher fraction of sites predicted to conform to SH2 binding rules. However, it was also an experiment to evaluate how much beyond wild type rules sSH2 domains can sample. We found that the superbinders did expand the sequence range as evidenced by the motif differences. The shift in motifs of sSH2 predicted negatives is a +1 acidic residue in the ligand, which is one of the key structural features that led to this work – one residue in the SH2 domain must contact both the negatively charged pY residue and the residue in the +1 position [19] – a match that can likely be satisfied by a negative charge in the +1 position. However, sSH2 enriched peptides still generally conform to the rules of SH2 recognition as evidenced by the evaluation of the predicted negative set, which still showed higher match to the rules of binding (higher overall prediction probabilities, compared to the negative set of pY-antibody enriched fractions). Hence, even with the expansion of the sSH2 recognition, superbinders are only capable of sampling the SH2 interaction space. This is a key consideration to be made in the use of these in proteomics pipelines. They represent an exciting new methodology for sampling deeper into the pY proteome and with newer pipelines may become even cheaper and more tractable for more labs to explore this area of proximal signaling space [7]. However, a complete shift to sSH2 alone will likely reduce the diversity of pY functions captured, regardless of what sSH2 domain (or combination) is used, since all SH2 domains share these global sequence rules for guiding interactions.

We believe that the general approach and the lessons learned in this work might be useful for larger generalizability across domain-motif interactions. Globular domains and short linear sequence interactions guide a large degree of protein interactions, from those with modifications (e.g. acetylation and bromodomains and phosphoserine and 14-3-3 or WW domains) and without modifications (e.g. SH3-polyproline interactions, kinase-substrates, and PDZ interactions with protein C-terminals). Hence, the combination of structural conservation analysis for evaluation of shared biophysical binding rules, coupled with machine learning to predict possible partners across the proteome might be highly useful across a broad range of interaction types. Additionally, we found that a relatively small amount of representative data was needed to build a sensitive and specific classifier, making this tractable as well for expansion. It is likely that data gathered from complementary experimental approaches and from diverse representatives of the family will be useful for new domain type classification, as it was for SH2 domains.

## Materials and Methods

### Peptide encoding

pY peptides were encoded using a combination of sequence-derived statistical and physiochemical features that captures positional preferences and biochemical properties relevant for binding.

#### i. DPPS-based encoding

Amino acids were represented using Divided Physicochemical Property Scores (DPPS) [49], in which each residue is encoded as a 10-dimensional vector describing its physicochemical properties. Position-specific DPPS profiles were constructed independently from the binder and non-binder training peptide sets. For each class *c* ∈ {B, NB}, where B and NB denote binder and nonbinder peptides, respectively, the position-specific DPPS profile was represented as a 6 × 10 matrix:

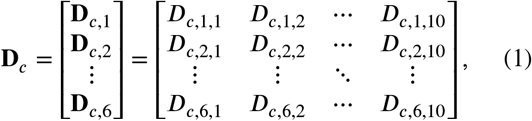

where each row represents the mean DPPS vector at a specific peptide position. The DPPS vector at position *j* was calculated as:

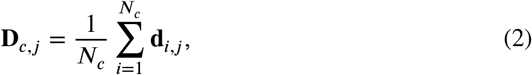

where **d**_***i***,*j*_ represents the 10-dimensional DPPS vector of the residue at position *j* in peptide ***i***, and *N*_*c*_ represents the number of peptides in class *c*. This procedure generated separate Binder (**D**_*B*_) and Nonbinder (**D**_*NB*_) DPPS profiles, each containing six position-specific 10-dimensional vectors.

To encode a query peptide, each residue was represented by its corresponding 10-dimensional DPPS vector, generating a query DPPS matrix:

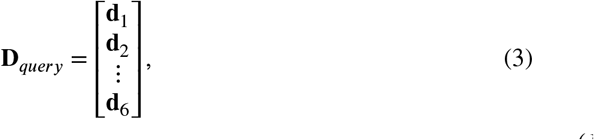

where **d**_*j*_ represents the DPPS vector of the residue at position *j*. The similarity of the query peptide to each class-specific DPPS profile was calculated by taking the dot product between corresponding positional DPPS vectors and summing across all six positions:

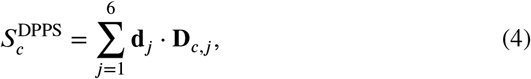

where **D**_*c,j*_ represents the position-specific DPPS vector from either the Binder (*c* = _*B*_) or Nonbinder (*c* = *N*_*B*_) profile. The resulting scores, 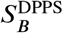 and 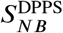, represent the DPPS-Binder and DPPS-Nonbinder features used for model training.

#### ii. Hydropathy-based encoding

Residue-level hydropathy values derived from the Kyte– Doolittle hydrophobicity scale [23] were used to encode peptide sequences. Each amino acid was represented by a single scalar value describing its hydrophobicity.

Position-specific hydropathy profiles were constructed independently from the binder and nonbinder training peptide sets. For each class *c* ∈ {B, NB}, the profile was represented as a six-dimensional vector containing the mean hydropathy value at each peptide position:

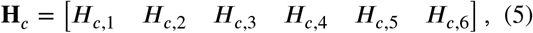

where each position-specific value was calculated as:

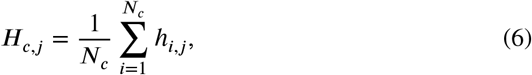

where *h*_***i***,*j*_ represents the hydropathy value of the residue at position *j* in peptide ***i***, and *N*_*c*_ represents the number of peptides in class *c*. This procedure generated separate Binder (**H**_*B*_) and Nonbinder (**H**_*NB*_) hydropathy profiles, each containing six position-specific hydropathy values.

To encode a query peptide, the hydropathy values of its six residues were represented as:

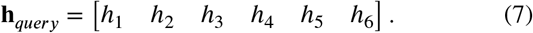

The similarity of the query peptide to the Binder and Nonbinder hydropathy profiles was calculated using a dot product between the query hydropathy vector and each class-specific profile:

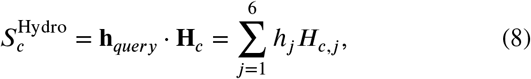

where *c* represents either the Binder (_*B*_) or Nonbinder (*N*_*B*_) class. The resulting scores, 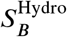 and 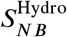, represent the Hydropathy-Binder and Hydropathy-Nonbinder features used for model training.

#### iii. Log-Odd positional scores

Position-specific Log-odds matrices (PSLMs) were constructed independently from the binder and nonbinder training peptide sets to capture amino acid preferences at each sequence position. For each class *c* ∈ {B, NB}, amino acid occurrences were counted independently at each of the six peptide positions to generate a position frequency matrix. A pseudocount of α = 0.5 was added to each amino acid count prior to normalization to avoid zero probabilities for amino acids not observed at a given position. The resulting probabilities were normalized such that the probabilities across the 20 amino acids at each position summed to one:

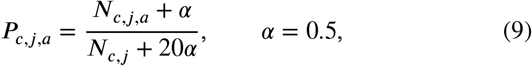

where *N*_*c,j,a*_ is the number of peptides in class *c* containing amino acid _*a*_ at position *j*, and *N*_*c,j*_ is the total number of peptides in class *c* at position *j*. The resulting probabilities were converted to log-odds scores relative to a uniform amino acid background distribution, *P*_bg,*a*_ = 1/20, as:

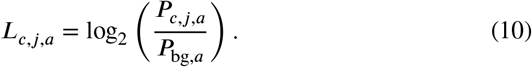

The resulting log-odds profile was represented as a 6×20 matrix:

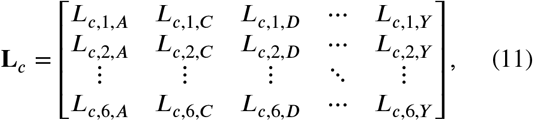

where each matrix element represents the log-odds score amino acid at peptide position *j*. This procedure genated separate Binder (**L**_*B*_) and Nonbinder (**L**_*NB*_) log-odds matrices.

To encode a query peptide, each residue was assigned corresponding log-odds score from the Binder and Non-binder profiles based on its amino acid identity and sequence position. For each class, the six residue-level log-odds values were summed across the peptide sequence to generate a class-specific log-odds score:

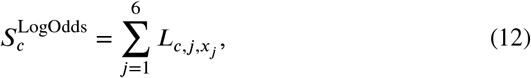

where *x*_*j*_ represents the amino acid at position *j* of the query peptide. The resulting scores, 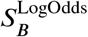 and 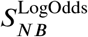, represent the LogOdds-Binder and LogOdds-Nonbinder features, respectively.

#### Feature construction and scaling

The three encoding strategies (DPPS, hydropathy, and log-odds) each generated two class-specific similarity scores corresponding to the binder and nonbinder profiles. For each six-residue peptide, these scores were concatenated to generate a six-dimensional feature representation:

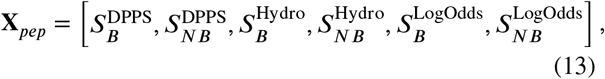

where each feature represents the similarity of the peptide sequence to either the binder (_*B*_) or nonbinder (*N*_*B*_) sequence-derived profile for the corresponding encoding strategy.

Feature scaling was performed using min–max normalization. Scaling parameters were fitted exclusively on the training data within each cross-validation fold and applied to the corresponding held-out test set without refitting, ensuring consistent feature transformation and preventing information leakage.

### SpY-C model Construction, Optimization, and Deployment

#### Support vector machine classifier

A support vector machine (SVM) classifier with a radial basis function (RBF) kernel was used for peptide classification. To account for class imbalance between binder and nonbinder peptides, the class weight parameter was set to “balanced”, which adjusts class contributions during model training according to their relative frequencies.

#### Model prototyping and selection using nested cross-validation

During model development, for each candidate training dataset, model performance was assessed using nested stratified cross-validation consisting of an outer evaluation loop and an inner hyperparameter optimization loop. In each outer split, the training portion was passed to a five-fold inner cross-validation, while the outer test fold remained held out from model optimization. Within each inner fold, sequence-derived feature profiles were generated exclusively from the inner-training peptides and subsequently used to encode the corresponding validation peptides. An RBF-kernel SVM was then evaluated across combinations of *C* ∈ {0.01, 0.1, 0.5, 1, 2, 5, 10} and γ ∈ {0.001, 0.01, 0.05, 0.1, 1}, with MinMax scaling fitted independently within each training fold. The hyperparameter combination yielding the highest mean ROC-AUC across the five inner validation folds was selected. Using these parameters, feature profiles were rebuilt from the complete outer-training set, the model was refitted, and performance was evaluated on the held-out outer test set. The outer cross-validation consisted of five folds repeated twice, and performance was summarized across the resulting test folds using ROC-AUC, accuracy, and F1 score. This step generated a nested cross-validation performance file containing fold-specific accuracy, F1 score, ROC-AUC, selected C and γ values, and inner cross-validation ROC-AUC across the 10 outer-fold evaluations.

### Final deployment model

Following nested cross-validation, stable hyperparameters for the final SpY-C deployment model were selected using repeated stratified cross-validation on the complete optimized training dataset containing all labeled binder and nonbinder 6-mer pY peptides. Ten independent five-fold cross-validation runs were performed, with the fold assignments varied between runs. Within each fold, sequence-derived feature profiles were generated exclusively from the training peptides and used to encode the corresponding validation peptides, thereby preventing information leakage during hyperparameter selection. All combinations of C and γ were evaluated using an RBF-kernel SVM with MinMax feature scaling and balanced class weights, and ROC-AUC was calculated for each validation fold.

For each *C* and γ combination, ROC-AUC values were first averaged across the five folds within each run and then summarized across the 10 runs to obtain the mean and standard deviation of the run-level AUCs. The hyperparameter combinations with the highest mean ROC-AUC across runs was selected. Candidates within 10™4 AUC of the highest-performing combination were considered equivalent, with lower C, followed by lower γ, used as tie-breakers. The sequence-derived feature profiles were then rebuilt using the complete training dataset, and the final RBF-SVM was trained once on all available training peptides using the selected hyperparameters.

The final SpY-C model was saved as a serialized .pkl artifact containing the trained SVM and fitted MinMax scaler, together with the frozen binder and nonbinder log-odds matrices, DPPS profiles, and hydropathy profiles required for peptide encoding. The artifact also retained the selected hyperparameters and model-selection and nested cross-validation performance statistics. For subsequent prediction tasks, this artifact was loaded directly without additional model training or parameter optimization. A separate candidate-selection file containing the mean and standard deviation of ROC-AUC across the 10 runs for each C and γ combination was also generated.

### Estimating Binder proportions using subsampling

To assess the robustness of predictions of peptides to variations in dataset composition, we performed a subsampling-based sensitivity analysis. For each dataset, the predicted binder proportion was calculated as the fraction of peptides classified as binders by SpY-C. Dataset stability was evaluated through repeated random subsampling without replacement, where 95% of peptides were randomly selected in each iteration and the corresponding binder fraction was recalculated. This procedure was repeated 10,000 times per dataset to generate a distribution of binder fraction estimates. The mean and standard deviation of these distributions were used to quantify the variability of predicted binder proportions under small changes in dataset composition. Lower variability indicates that the estimated binder fraction is robust to peptide-level perturbations, whereas higher variability suggests greater dependence on specific peptides within the dataset.

### Bootstrap distributions of median SH2-binding probabilities

To compare SpY-C prediction probabilities across peptide populations while accounting for differences in dataset size, we randomly sampled 50 unique peptides without replacement from each population and calculated the median prediction probability. This process was repeated 5,000 times to generate distributions of median prediction probabilities. The resulting distributions were visualized using kernel density estimates (KDEs).

### ClinVar variant mapping to phosphotyrosine sites

The ClinVar variant summary file variant_summary.txt.gz was downloaded from the NCBI FTP repository [24], and variants corresponding to human genes of interest were extracted based on UniProt gene mappings. Protein-level HGVS annotations were parsed from ClinVar variant names to identify missense substitutions and obtain the wild-type residue, amino acid position, and substituted residue. These variants were subsequently mapped to known phosphotyrosine sites in the human phosphoproteome to identify substitutions occurring within the SH2-binding sequence window (pY-2 to pY+4) surrounding each pY site for downstream SpY-C analysis.

### Mutation enrichment analysis

To identify sequence changes associated with altered SH2-binding potential, residue- and position-specific enrichment analyses were performed separately for predicted gain- and loss-of-function mutations. For each category, the foreground consisted of mutations classified as gain- or loss-of-function, while the background comprised the remaining wild-type–mutant pairs originating from the same initial prediction class. Enrichment of individual amino acid substitutions at each position within the SH2-binding window was assessed using Fisher’s exact test. *P* values were corrected for multiple testing using the Benjamini–Hochberg procedure, and odds ratios were used to quantify enrichment. Substitutions with an FDR-adjusted *P* < 0.05 and odds ratio > 1 were considered significantly enriched.

### ProteomeScout-based peptide preparation and annotation

Datasets requiring biological annotation of phosphotyrosine-containing peptides were processed using ProteomeScout.

Dataset preparation was performed in July 2026 using the ProteomeScout web interface (version 4), ProteomeScoutAPI version 3.1.1, and ProteomeScout data version 8. Peptide sequences were formatted according to ProteomeScout requirements, and protein identifiers were standardized using UniProt identifiers. The datasets subjected to this annotation procedure are indicated in (https://doi.org/10.6084/m9.figshare.33256416).

### SH2 Superbinder pull downs and mass spectrometry

Jurkat PD-1+_GFP T cells (cells expressing PD-1 receptor conjugated to GFP) were stimulated by using pre-conjugated CD3 and CD28 antibodies with Dynabeads Protein G magnetic beads for 2 minutes. Non-stimulated cells were performed by adding empty Dynabeads Protein G magnetic beads instead. After incubation, cell stimulation was stopped by addition of equal volume of hot 8M Guanidine-HCl in 100 mM HEPES pH 8.0 (diluting it to 4M Guan-HCl final) and heated for 5min at 95°C, followed by sonication for 30 seconds. The magnetic beads were removed using a magnet and the samples were then subjected to an additional centrifugation step at 14,000rpm for 15min at 4°C to remove residual debris. PierceTM BCA Protein Assay Kit (Thermo Fisher Scientific) was used to measure protein concentrations.

Protein extracts were then reduced with 1 mM TCEP (Tris(2-carboxyethyl)phosphine hydrochloride) for 30 min at room temperature (RT), alkylated with 11 mM CAA (chloroacetamide) for 30 min at RT in the dark. Samples were then diluted with 50 mM HEPES buffer pH 8.0 to 1M Guanidine-HCl and digested with Lys-C (Wako) at a 1:100 enzyme/protein (w/w) ratio for 10 hours at RT, followed by digestion with 1:100 trypsin (Promega) overnight at RT. Subsequently, samples were acidified (pH 2.0–3.0) using TFA (trifluoroacetic acid), desalted using Sep-Pak C18 classic cartridges (Waters), peptide concentration measured by nanodrop and samples were lyophilized ON, all steps as previously described [1]. Lyophilized samples were dissolved in ice cold IAP buffer (50mM MOPS pH 7.6; 10mM Na_2_HPO_4_; 50mM NaCl) containing 0.1% (v/v) Triton-X-100. For each pull down, 1000*µ*g tryptic peptides was incubated with 50*µ*L slurry of SH2 Superbinder agarose beads (Precision Proteomics) for 4h at 4°C in slow agitation (10 rpm). The beads were then washed 8 times in ice cold IAP buffer followed by a single wash with 150 mM NaCl. Phosphopeptides were eluted sequentially by 5 min incubation in 0.3% (v/v) TFA once and 5min incubation in 0.1% (v/v) TFA twice.

Peptide samples were desalted, concentrated and analyzed on an LC-MS/MS system consisting of an EASY-nLC 1000 nanoflow liquid chromatograph coupled with an Orbitrap Exploris 480 mass spectrometer (Thermo Fisher Scientific) using DIA acquisition method as described [5]. Briefly, for MS1, we used a 350 to 1400m/z survey scan with a resolution of 120,000, a maximum ion injection time of 45ms, and a normalized automatic gain control target of 300%. This was followed by MS2 fragmentation of precursor ions by higher-collisional dissociation at a collision energy of 28% for each 13m/z isolation window, and a 1m/z overlap between sliding windows to sequentially cover the 361-1000m/z range by 50 scans. For MS2, resolution was set to 30,000 and the maximum ion injection time to 54ms. DIA raw files were analyzed in SpectronautTM version 18.1 (Biognosys) using default settings for Phospho PTM workflow. The report table containing phosphopeptide identifications and quantitative values were kept for subsequent data analysis in R. The mass spectrometry proteomics data have been deposited to the ProteomeXchange Consortium via the PRIDE partner repository with the dataset identifier PXD079717 and https://doi.org/doi:10.6019/PXD079717. The processed PRIDE dataset used in this study is provided as a supplementary data file (Supplemental_Data_1.xlsx).

### PepspotDB (SPOT Array Data)

The SPOT Array dataset used in this study that is no longer fully accessible through their original sources. We have recompiled, processed, and deposited on Figshare (https://doi.org/10.6084/m9.figshare.33249876), along with processing details, to ensure continued availability and reproducibility. This resource includes the original experiment files detailing each SH2 probe to a single SPOT array and includes the signal, p-value calculated according to the original study authors, the peptide sequence and the neural network prediction value as part of that original study [50]. We updated the protein references for peptides and SH2 domains to current Uniprot identifiers (from original NCBI GI accessions). Phosphopeptides used in the original study included phosphotyrosines known at that time, along with tyrosines in the human proteome predicted to be phosphorylated. We used ProteomeScout web version 4 (https://proteomescout.research.virginia.edu/), running ProteomeScoutAPI v3.1.1 and ProteomeScout data version 7 to annotate the peptides – aligning peptides with current Uniprot records and annotating whether the tyrosine is known to be modified currently. Finally, we manually updated some SH2 domain references and verified their match to current protein records and included the species sequence used at that time. In the Figshare resource for modernized PepspotDB, we also provide a reshaped signal matrix, which provides the signal intensity values for each peptide (rows) and the experimental SH2 (column), along with the annotation of whether the phosphorylation site is found and we used the original experimental quality value to provide no value in a column if the quality was indicated as “BAD” in the original experimental files. Given the signal intensity data for SH2 domains did not conform to normal distributions, when we reshaped the data, we selected to ignore the p-value, which was calculated based on standard deviations from the mean, and kept all signal intensity values. Unless noted, in SpY-C, we used all peptides in the top 5% as hits for each domain, not selecting for annotated phosphotyrosines in the current proteome. For the analyses presented in this work, only peptides classified as having a good alignment status were included and SH2 domain sequences from Homo sapiens were included. We found excellent concordance between replicates, where multiple experimental replicates were available for an SH2 domain so, for this work, we used the first replicate of any duplicated experiments.

### Dataset collection and processing

The datasets analyzed in this study were collected from multiple published sources and organized into four major categories: (i) experimentally measured SH2–pY binding datasets; (ii) human and mouse proteome sequences; (iii) phosphoproteomic datasets from large-scale mass spectrometry studies, tyrosine-dependent signaling stimulation experiments, and affinity-enrichment experiments; and (iv) ClinVar missense mutations. For each dataset, the published information was curated and processed to retain the sequence and experimental information necessary to identify the corresponding peptide motifs and the conditions under which they were identified in the original study. The processed datasets used for analysis, together with the resulting SpY-C predictions, are provided through Figshare (https://doi.org/10.6084/m9.figshare.33256416).

## Code availability and model reproducibility

The SpY-C codebase used for model training, evaluation, and deployment is publicly available through the GitHub repository (https://github.com/NaegleLab/SpY-C). Reproducibility of the deployed model (SpY-C_v1.0) is ensured by providing the final trained model artifact (.pkl file) generated using the final training dataset and selected hyperparameters. Loading this artifact and applying it to the same peptide sequences using the provided deployment code produces identical predictions to those reported in this study. Instructions for loading the model artifact and reproducing the predictions are provided in the GitHub repository. The GitHub codebase used for this work is archived in Figshare and is available at (https://doi.org/10.6084/m9.figshare.33256800).

## Acknowledgements

Research reported in this publication was supported by the National Institute Of General Medical Sciences of the National Institutes of Health under Award Number R35GM138127 and the National Institute of Allergy and Infectious Disease under Award Number R01AI153617. The content is solely the responsibility of the authors and does not necessarily represent the official views of the National Institutes of Health. We would like to thank Dr. Gianni Cesarani for allowing us to use and distribute the PepspotDB study.

## Bibliography References

[1] Akimov, V., Barrio-Hernandez, I., Hansen, S.V., Hallenborg, P., Pedersen, A.K., Bekker-Jensen, D.B., Puglia, M., Christensen, S.D., Vanselow, J.T., Nielsen, M.M., et al., 2018. Ubisite approach for comprehensive mapping of lysine and n-terminal ubiquitination sites. Nature structural & molecular biology 25, 631–640.

[2] Batth, T.S., Papetti, M., Pfeiffer, A., Tollenaere, M.A., Francavilla, C., Olsen, J.V., 2018. Large-scale phosphoproteomics reveals shp-2 phosphatase-dependent regulators of pdgf receptor signaling. Cell reports 22, 2784–2796.

[3] Beebe, K.D., Wang, P., Arabaci, G., Pei, D., 2000. Determination of the binding specificity of the sh2 domains of protein tyrosine phosphatase shp-1 through the screening of a combinatorial phosphotyrosyl peptide library. Biochemistry 39, 13251–13260.

[4] Bian, Y., Li, L., Dong, M., Liu, X., Kaneko, T., Cheng, K., Liu, H., Voss, C., Cao, X., Wang, Y., et al., 2016. Ultra-deep tyrosine phosphoproteomics enabled by a phosphotyrosine superbinder. Nature chemical biology 12, 959–966.

[5] Boel, F., Akimov, V., Teuchler, M., Terkelsen, M.K., Wernberg, C.W.,Larsen, F.T., Hallenborg, P., Lauridsen, M.M., Krag, A., Mandrup, S., et al., 2025. Deep proteome profiling of metabolic dysfunction-associated steatotic liver disease. Communications medicine 5, 56.

[6] Chang, A., Leutert, M., Rodriguez-Mias, R.A., Villén, J., 2023. Automated enrichment of phosphotyrosine peptides for high-throughput proteomics. bioRxiv.

[7] Chang, A.T., Rodriguez-Mias, R.A., Berg, M.D., Moggridge, S.,Villén, J., 2026. Scalable phosphotyrosine enrichment with SH2 superbinder enables deep profiling of EGF responses. The EMBO Journal 45, 5556–5579. URL: https://doi.org/10.1038/s44318-026-00843-8, doi:10.1038/s44318-026-00843-8.

[8] Chylek, L.A., Akimov, V., Dengjel, J., Rigbolt, K.T., Hu, B.,Hlavacek, W.S., Blagoev, B., 2014. Phosphorylation site dynamics of early t-cell receptor signaling. PloS one 9, e104240.

[9] Cochrane, D., Webster, C., Masih, G., McCafferty, J., 2000. Identification of natural ligands for sh2 domains from a phage display cdna library. Journal of Molecular Biology 297, 89–97.

[10] Doytchinova, I.A., Blythe, M.J., Flower, D.R., 2002. Additive method for the prediction of protein-peptide binding affinity. application to the mhc class i molecule hla-a* 0201. Journal of proteome research 1, 263–272.

[11] Gagoski, D., Rube, H.T., Rastogi, C., Melo, L.A., Li, X., Voleti, R., Shah, N.H., Bussemaker, H.J., 2025. Accurate sequence-to-affinity models for sh2 domains from multi-round peptide binding assays coupled with free-energy regression. bioRxiv, 2024–12.

[12] Gill, K., Macdonald-Obermann, J.L., Pike, L.J., 2017. Epidermal growth factor receptors containing a single tyrosine in their c-terminal tail bind different effector molecules and are signaling-competent. Journal of Biological Chemistry 292, 20744–20755.

[13] Henriques, D.A., Ladbury, J.E., Jackson, R.M., 2000. Comparison of binding energies of srcsh2-phosphotyrosyl peptides with structure-based prediction using surface area based empirical parameterization. Protein Science 9, 1975–1985.

[14] Hu, J., Liu, J., Ghirlando, R., Saltiel, A.R., Hubbard, S.R., 2003.Structural basis for recruitment of the adaptor protein aps to the activated insulin receptor. Molecular cell 12, 1379–1389.

[15] Huang, H., Li, L., Wu, C., Schibli, D., Colwill, K., Ma, S., Li, C., Roy, P., Ho, K., Songyang, Z., et al., 2008. Defining the specificity space of the human src homology 2 domain. Molecular & Cellular Proteomics 7, 768–784.

[16] Hunter, T., 2000. Signaling—2000 and beyond. Cell 100, 113–127.

[17] Jin, L.L., Wybenga-Groot, L.E., Tong, J., Taylor, P., Minden, M.D., Trudel, S., McGlade, C.J., Moran, M.F., 2015. Tyrosine phosphorylation of the lyn src homology 2 (sh2) domain modulates its binding affinity and specificity*[s]. Molecular & Cellular Proteomics 14, 695–706.

[18] Jones, R.B., Gordus, A., Krall, J.A., MacBeath, G., 2006. A quantitative protein interaction network for the erbb receptors using protein microarrays. Nature 439, 168–174.

[19] Kandoor, A., Martinez, G., Hitchcock, J.M., Angel, S., Campbell, L., Rizvi, S., Naegle, K.M., 2025. Codiac: A comprehensive approach for interaction analysis provides insights into sh2 domain function and regulation. Science signaling 18, eads8396.

[20] Kaneko, T., Huang, H., Cao, X., Li, X., Li, C., Voss, C., Sidhu, S.S., Li, S.S., 2012. Superbinder sh2 domains act as antagonists of cell signaling. Science signaling 5, ra68–ra68.

[21] Kundu, K., Costa, F., Huber, M., Reth, M., Backofen, R., 2013. Semi-supervised prediction of sh2-peptide interactions from imbalanced high-throughput data. PloS one 8, e62732.

[22] Kundu, K., Mann, M., Costa, F., Backofen, R., 2014. Modpepint:an interactive web server for prediction of modular domain–peptide interactions. Bioinformatics 30, 2668–2669.

[23] Kyte, J., Doolittle, R.F., 1982. A simple method for displaying the hydropathic character of a protein. Journal of molecular biology 157, 105–132.

[24] Landrum, M.J., Lee, J.M., Riley, G.R., Jang, W., Rubinstein, W.S.,Church, D.M., Maglott, D.R., 2014. Clinvar: public archive of relationships among sequence variation and human phenotype. Nucleic acids research 42, D980–D985.

[25] Lee, J.K., Moon, T., Chi, M.W., Song, J.S., Choi, Y.S., Yoon, C.N., 2003. An investigation of phosphopeptide binding to sh2 domain. Biochemical and biophysical research communications 306, 225–230.

[26] Li, A., Voleti, R., Lee, M., Gagoski, D., Shah, N.H., 2023. High-throughput profiling of sequence recognition by tyrosine kinases and sh2 domains using bacterial peptide display. Elife 12, e82345.

[27] Li, L., Wu, C., Huang, H., Zhang, K., Gan, J., Li, S.S.C., 2008. Prediction of phosphotyrosine signaling networks using a scoring matrix-assisted ligand identification approach. Nucleic acids research 36, 3263–3273.

[28] Liu, B.A., Engelmann, B.W., Nash, P.D., 2012. The language of sh2domain interactions defines phosphotyrosine-mediated signal transduction. FEBS letters 586, 2597–2605.

[29] Liu, B.A., Jablonowski, K., Raina, M., Arcé, M., Pawson, T., Nash, P.D., 2006. The human and mouse complement of sh2 domain proteins—establishing the boundaries of phosphotyrosine signaling. Molecular cell 22, 851–868.

[30] Liu, B.A., Jablonowski, K., Shah, E.E., Engelmann, B.W., Jones,R.B., Nash, P.D., 2010. Sh2 domains recognize contextual peptide sequence information to determine selectivity. Molecular & Cellular Proteomics 9, 2391–2404.

[31] Liu, H., Li, L., Voss, C., Wang, F., Liu, J., Li, S.S.C., 2015. A comprehensive immunoreceptor phosphotyrosine-based signaling network revealed by reciprocal protein–peptide array screening. Molecular & Cellular Proteomics 14, 1846–1858.

[32] Martyn, G.D., Veggiani, G., Kusebauch, U., Morrone, S.R., Yates,B.P., Singer, A.U., Tong, J., Manczyk, N., Gish, G., Sun, Z., et al., 2022. Engineered sh2 domains for targeted phosphoproteomics. ACS chemical biology 17, 1472.

[33] Matlock, M.K., Holehouse, A.S., Naegle, K.M., 2015. Proteomescout: a repository and analysis resource for post-translational modifications and proteins. Nucleic acids research 43, D521–D530.

[34] Moran, M.F., Koch, C.A., Anderson, D., Ellis, C., England, L.,Martin, G.S., Pawson, T., 1990. Src homology region 2 domains direct protein-protein interactions in signal transduction. Proceedings of the National Academy of Sciences 87, 8622–8626.

[35] Needham, E.J., Parker, B.L., Burykin, T., James, D.E., Humphrey, S.J., 2019. Illuminating the dark phosphoproteome. Science signaling 12, 1–18.

[36] Obenauer, J.C., Cantley, L.C., Yaffe, M.B., 2003. Scansite 2.0:Proteome-wide prediction of cell signaling interactions using short sequence motifs. Nucleic acids research 31, 3635–3641.

[37] Ochoa, D., Jarnuczak, A.F., Viéitez, C., Gehre, M., Soucheray, M., Mateus, A., Kleefeldt, A.A., Hill, A., Garcia-Alonso, L., Stein, F., Krogan, N.J., Savitski, M.M., Swaney, D.L., Vizcaíno, J.A., Noh, K.M., Beltrao, P., 2020. The functional landscape of the human phosphoproteome. Nature Biotechnology 38, 365–373. URL: https://doi.org/10.1038/s41587-019-0344-3, doi:10.1038/s41587-019-0344-3.

[38] Pawson, T., 2004. Specificity in signal transduction: from phosphotyrosine-sh2 domain interactions to complex cellular systems. Cell 116, 191–203.

[39] Pawson, T., Gish, G.D., 1992. Sh2 and sh3 domains: from structure to function. Cell 71, 359–362.

[40] Pawson, T., Gish, G.D., Nash, P., 2001. Sh2 domains, interaction modules and cellular wiring. Trends in cell biology 11, 504–511.

[41] Pawson, T., Nash, P., 2000. Protein–protein interactions define specificity in signal transduction. Genes & development 14, 1027–1047.

[42] Rahuel, J., Gay, B., Erdmann, D., Strauss, A., García-Echeverría, C., Furet, P., Caravatti, G., Fretz, H., Schoepfer, J., Grütter, M.G., 1996. Structural basis for specificity of grb2-sh2 revealed by a novel ligand binding mode. Nature structural biology 3, 586–589.

[43] Rodriguez, M., Li, S.S.C., Harper, J.W., Songyang, Z., 2004. An oriented peptide array library (opal) strategy to study protein-protein interactions. Journal of Biological Chemistry 279, 8802–8807.

[44] Ronan, T., Garnett, R., Naegle, K.M., 2020. New analysis pipeline for high-throughput domain–peptide affinity experiments improves sh2 interaction data. Journal of Biological Chemistry 295, 11346–11363.

[45] Sánchez, I.E., Beltrao, P., Stricher, F., Schymkowitz, J., Ferkinghoff-Borg, J., Rousseau, F., Serrano, L., 2008. Genome-wide prediction of sh2 domain targets using structural information and the foldx algorithm. PLoS computational biology 4, e1000052.

[46] Songyang, Z., Shoelson, S., McGlade, J., Olivier, P., Pawson, T., Bustelo, X., Barbacid, M., Sabe, H., Hanafusa, H., Yi, T., et al., 1994. Specific motifs recognized by the sh2 domains of csk, 3bp2, fps/fes, grb-2, hcp, shc, syk, and vav. Molecular and cellular biology 14, 2777–2785.

[47] Suenaga, A., Ichikawa, M., Hatakeyama, M., Yu, X., Futatsugi, N., Narumi, T., Fukui, K., Terada, T., Taiji, M., Shirouzu, M., et al., 2003. Molecular dynamics, free energy, and spr analyses of the interactions between the sh2 domain of grb2 and erbb phosphotyrosyl peptides. Biochemistry 42, 5195–5200.

[48] Sundberg, E.J., Urrutia, M., Braden, B.C., Isern, J., Tsuchiya, D., Fields, B.A., Malchiodi, E.L., Tormo, J., Schwarz, F.P., Mariuzza, R.A., 2000. Estimation of the hydrophobic effect in an antigen-antibody protein-protein interface. Biochemistry 39, 15375–15387.

[49] Tian, F., Yang, L., Lv, F., Yang, Q., Zhou, P., 2009. In silico quantitative prediction of peptides binding affinity to human mhc molecule: an intuitive quantitative structure–activity relationship approach. Amino Acids 36, 535–554.

[50] Tinti, M., Kiemer, L., Costa, S., Miller, M.L., Sacco, F., Olsen, J.V., Carducci, M., Paoluzi, S., Langone, F., Workman, C.T., et al., 2013. The sh2 domain interaction landscape. Cell reports 3, 1293–1305.

[51] Videlock, E.J., Chung, V.K., Hall, J.M., Hines, J., Agapakis, C.M.,Austin, D.J., 2005. Identification of a molecular recognition role for the activation loop phosphotyrosine of the src tyrosine kinase. Journal of the American Chemical Society 127, 1600–1601.

[52] Wavreille, A.S., Garaud, M., Zhang, Y., Pei, D., 2007. Defining sh2 domain and ptp specificity by screening combinatorial peptide libraries. Methods 42, 207–219.

[53] Wolf-Yadlin, A., Hautaniemi, S., Lauffenburger, D.A., White, F.M.,2007. Multiple reaction monitoring for robust quantitative proteomic analysis of cellular signaling networks. Proceedings of the National Academy of Sciences 104, 5860–5865.

[54] Yaffe, M.B., Leparc, G.G., Lai, J., Obata, T., Volinia, S., Cantley, L.C., 2001. A motif-based profile scanning approach for genome-wide prediction of signaling pathways. Nature biotechnology 19, 348–353.

[55] Zhou, S., Shoelson, S.E., Chaudhuri, M., Gish, G., Pawson, T., Haser,W.G., King, F., Roberts, T., Ratnofsky, S., Lechleider, R.J., et al., 1993. Sh2 domains recognize specific phosphopeptide sequences. Cell 72, 767–778.

[56] Zhuang, G., Yu, K., Jiang, Z., Chung, A., Yao, J., Ha, C., Toy, K., Soriano, R., Haley, B., Blackwood, E., et al., 2013. Phosphoproteomic analysis implicates the mtorc2-foxo1 axis in vegf signaling and feedback activation of receptor tyrosine kinases. Science signaling 6, ra25–ra25.

