## Supplementary Figures for "SpY-C: Supervised Learning of Phosphopeptide Sequence Constraints Enables Global Prediction of SH2 Domain Binding"

### This file includes:

**Supplementary Figure S1** Projection of peptide representations into different feature spaces to evaluate separation between SH2 binders and nonbinders.

**Supplementary Figure S2** Evaluation of positive training set composition and optimization through additional SH2 domain sets.

**Supplementary Figure S3** Assessment of model generalizability using predicted binder proportions across known human phosphotyrosine sites.

**Supplementary Figure S4** Predicting binder proportions across engineered sSH2 domains.

**Supplementary Figure S5** Phosphotyrosine coverage and activation-loop peptide recovery by antibody and sSH2 enrichment.

**Supplementary Figure S6** Sequence motif profiles of peptides used for model training.

**Supplementary Figure S7** Sequence motif analysis of peptides enriched across antibody and superbinder datasets.

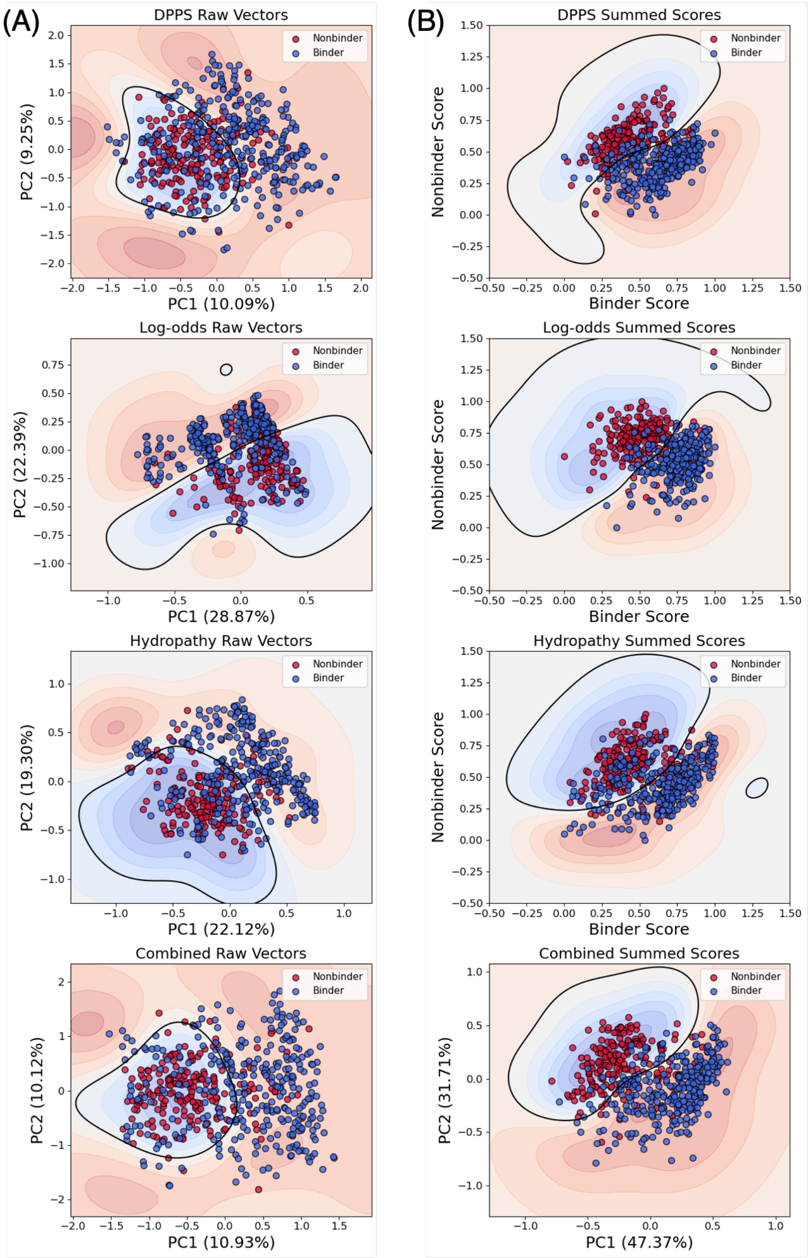


**Supplementary Figure S1.** Projection of peptide representations into different feature spaces to evaluate separation between SH2 binders and nonbinders. Each peptide (binder and nonbinder) was encoded using three sequence-derived feature representations: dipeptide physicochemical property scores (DPPS), position-specific log-odds scores, and hydropathy profiles. These encodings were evaluated in two forms: **(A)** raw feature vectors, where each peptide is represented by its full positional feature representation, and **(B)** aggregated profile-based scores, where each peptide is assigned a summed similarity score against binder and nonbinder reference profiles generated as described in Methods. In addition, combined representations integrating DPPS, log-odds, and hydropathy features were evaluated for both raw and aggregated feature spaces. Because raw feature representations are high-dimensional, principal component analysis (PCA) was used to project these vectors into two dimensions for visualization. Aggregated representations were directly visualized using the binder and nonbinder profile scores as the two dimensions. For each two-dimensional feature space, an RBF-kernel support vector machine (SVM) was fitted directly to the projected coordinates to visualize class separation, with the black contour line representing the SVM decision boundary. Binder and nonbinder peptides are shown in blue and red, respectively. These projections demonstrate the extent to which individual and combined sequence-derived feature representations capture the underlying sequence determinants distinguishing SH2-binding and nonbinding phosphotyrosine peptides.

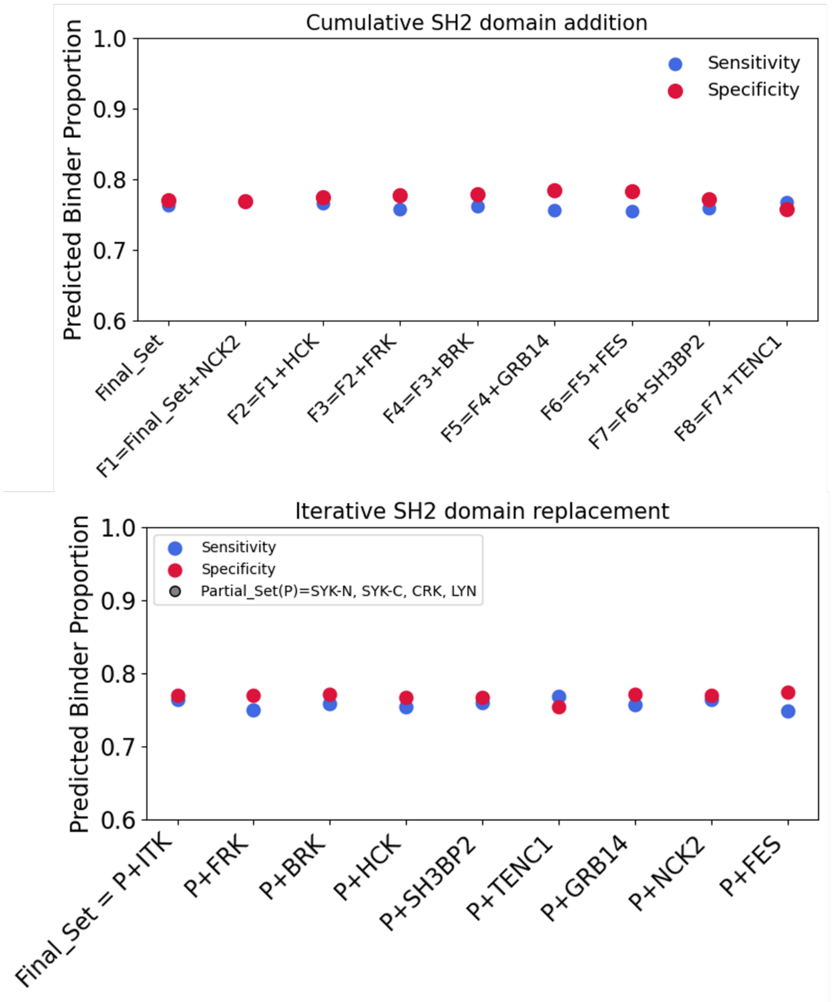


**Supplementary Figure S2.** Evaluation of positive training set composition and optimization through additional SH2 domain sets. To assess whether the composition of the final positive training set (represented as Final_Set) used for SpY-C development was optimized, we performed two complementary analyses. **(A)** SH2 domains were cumulatively incorporated to the 'Final_Set', and model sensitivity and specificity were evaluated after each addition. **(B)** To assess the robustness of the selected SH2 domain composition, approximately one-fifth of the positive training peptides from the 'Final_Set' were iteratively replaced with peptides derived from different SH2 domains shown to exhibit diverse binding specificities. The remaining four-fifths of the 'Final_Set', which were held constant across iterations, are referred to as the 'Partial_Set' (P). For each resulting training set combination, sensitivity and specificity were evaluated to determine whether model performance plateaued, with no further improvement upon incorporating or replacing additional SH2 domains.

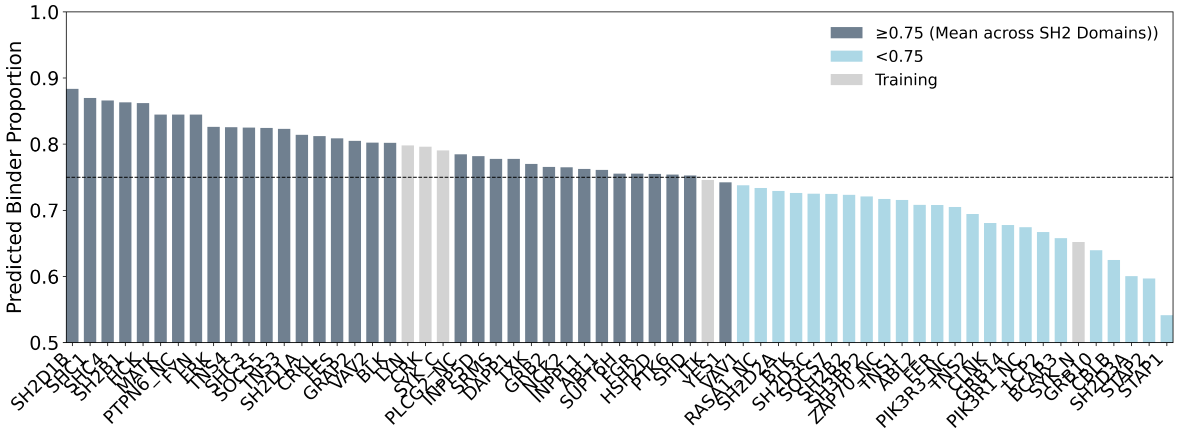


**Supplementary Figure S3.** Assessment of model generalizability using predicted binder proportions across known human phosphotyrosine sites. To assess model performance on physiologically relevant phosphotyrosine sequences, peptides evaluated for each SH2 domain were restricted to experimentally documented human phosphotyrosine (pY) sites from the human phosphoproteome using Proteomescout annotations. For each SH2 domain in the PepspotDB dataset, predicted binder proportions were calculated among peptides corresponding to the top 5% of experimentally observed enrichment signal intensities. The dotted horizontal line indicates the mean predicted binder proportion across all evaluated SH2 domains (0.75).

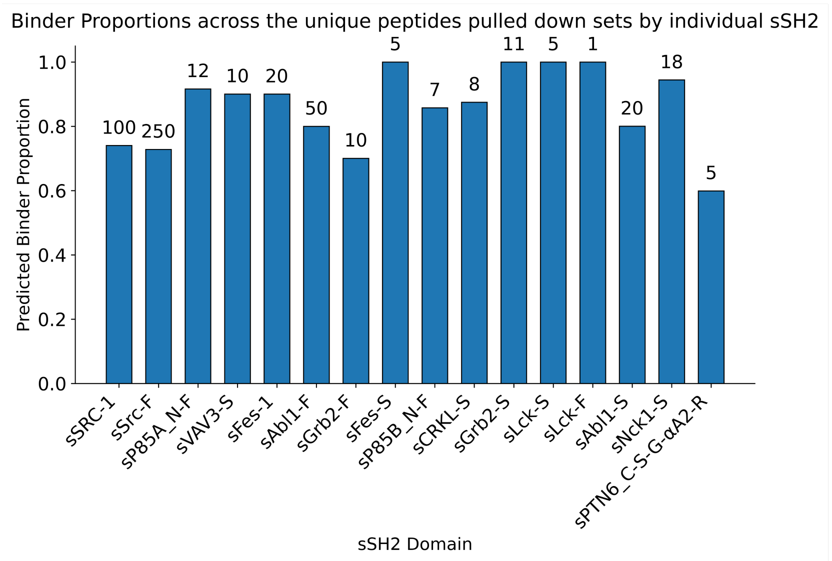


**Supplementary Figure S4.** Predicting binder proportions across engineered sSH2 domains. To evaluate whether the model captures differences in peptide recognition across engineered SH2 superbinder (sSH2) domains, peptides enriched by individual sSH2 domains from the AP-MS experiments reported by Martyn et al. were analyzed separately. For each sSH2 domain, we report the predicted binder proportion among the unique peptides enriched in each experimentally pulled-down peptide set. The number of unique peptides analyzed for each sSH2 domain is annotated above the corresponding bar.

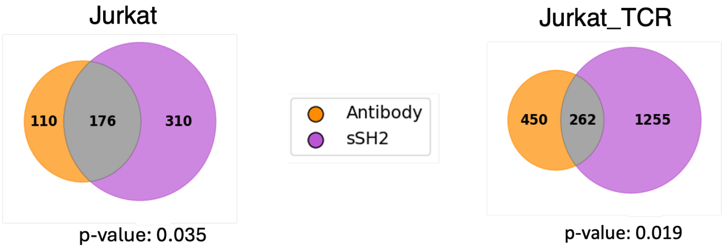


| | pY peptides from Activation loops | Other enriched pY peptides |
|---|---|---|
| Antibody | 7 | 279 |
| sSH2 | 3 | 483 |

| | pY peptides from Activation loops | Other enriched pY peptides |
|---|---|---|
| Antibody | 10 | 702 |
| sSH2 | 7 | 1510 |

**Supplementary Figure S5.** Phosphotyrosine coverage and activation-loop peptide recovery by antibody and sSH2 enrichment. **(A)** Venn diagrams show the overlap and unique pY peptides recovered by antibody- and sSH2-based enrichment across the indicated datasets. **(B)** Contingency tables summarize the recovery of pY peptides annotated within protein activation loops by each enrichment strategy. Depletion of activation-loop peptides in sSH2 relative to antibody enrichment was assessed using a one-sided Fisher's exact test, with odds ratios and P values indicated.

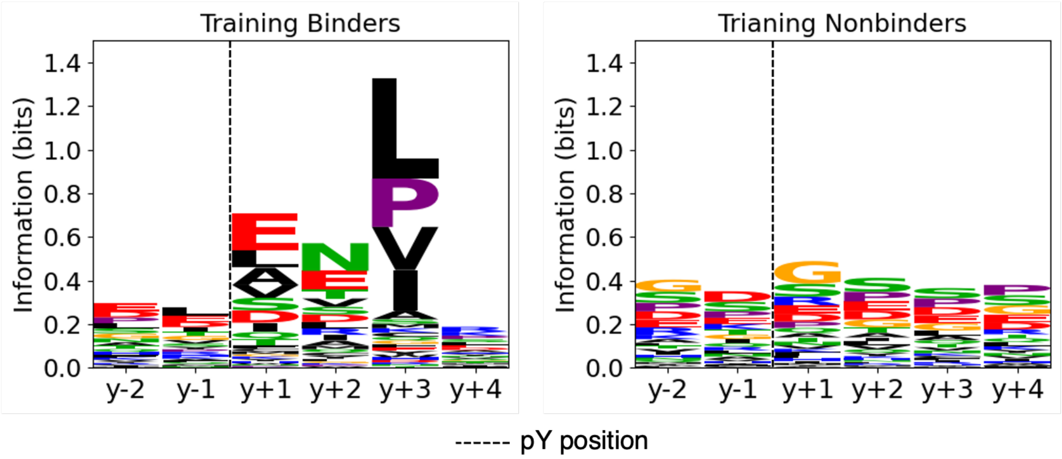


**Supplementary Figure S6.** Sequence motif profiles of peptides used for model training. Sequence motif logos were generated for the positive and negative training datasets used for final model SpY-C development. The position of the phosphotyrosine (pY) residue is indicated by a vertical dotted line.

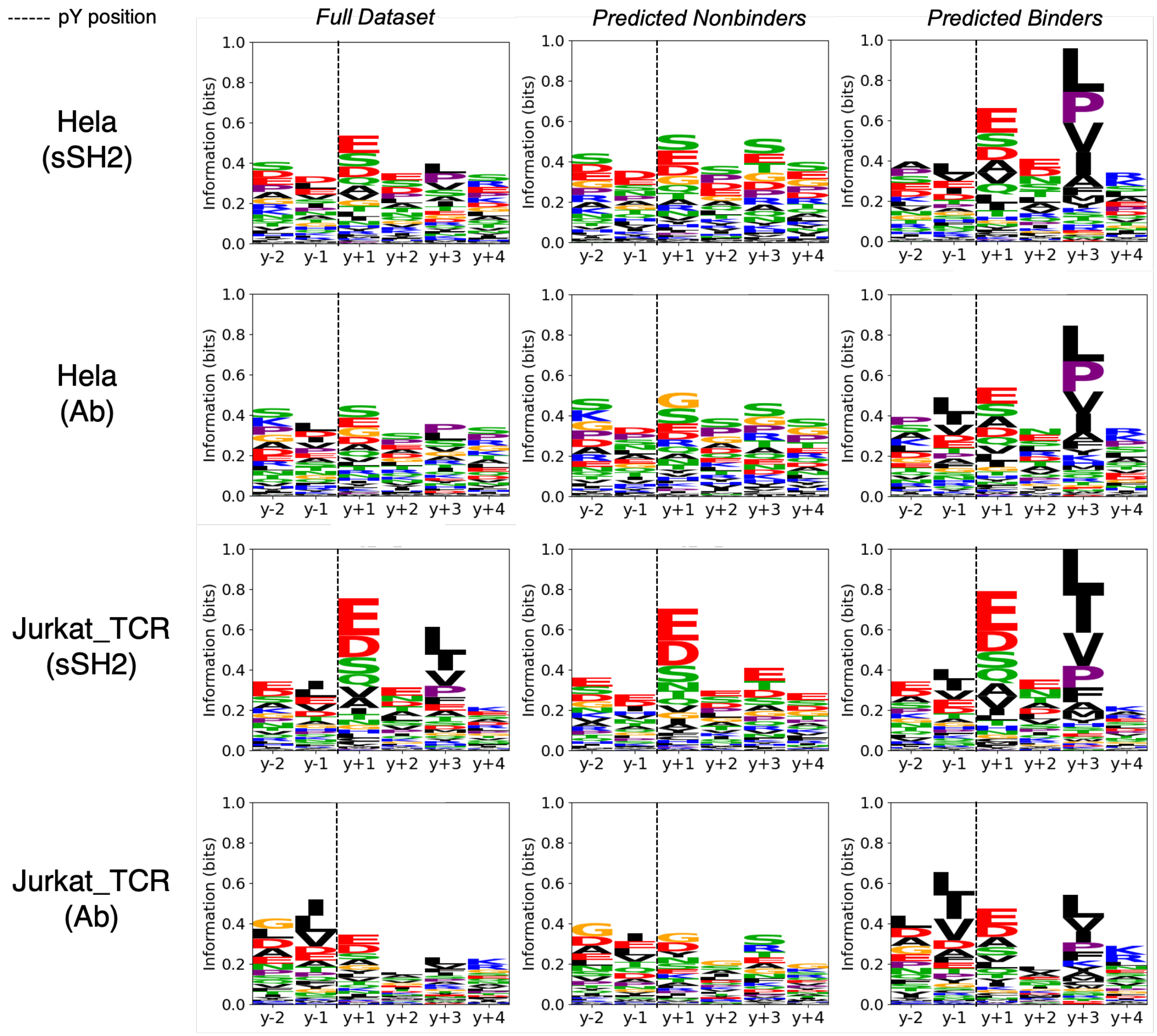


**Supplementary Figure S7.** Sequence motif analysis of peptides enriched across antibody and superbinder datasets. Sequence motif logos were generated to compare the peptide sequence profiles of phosphotyrosine peptides enriched by antibody and sSH2 superbinder strategies from the datasets analyzed in Figure 3A. For the Jurkat_TCR and HeLa datasets, motif logos were generated separately for the full peptide set used for prediction, as well as subsets classified as predicted binders and predicted nonbinders by SpY-C. Predicted nonbinder peptides, particularly from the Jurkat_TCR dataset, exhibit a pronounced enrichment of acidic residues at the +1 position. The position of the phosphotyrosine (pY) residue is indicated by a vertical dotted line.
